# Intragenomic rearrangements in SARS-CoV-2, other betacoronaviruses, and alphacoronaviruses

**DOI:** 10.1101/2022.03.07.483258

**Authors:** Roberto Patarca, William A. Haseltine

## Abstract

Variation of the betacoronavirus SARS-CoV-2 has been the bane of COVID-19 control. Documented variation includes point mutations, deletions, insertions, and recombination among closely or distantly related coronaviruses. Here, we describe yet another aspect of genome variation by beta- and alphacoronaviruses. Specifically, we report numerous genomic insertions of 5’-untranslated region sequences into coding regions of SARS-CoV-2, other betacoronaviruses, and alphacoronaviruses. To our knowledge this is the first systematic description of such insertions. In many cases, these insertions change viral protein sequences and further foster genomic flexibility and viral adaptability through insertion of transcription regulatory sequences in novel positions within the genome. Among human Embecorivus betacoronaviruses, for instance, from 65% to all of the surveyed sequences in publicly available databases contain 5’-UTR-derived inserted sequences. In limited instances, there is mounting evidence that these insertions alter the fundamental biological properties of mutant viruses. Intragenomic rearrangements add to our appreciation of how variants of SARS-CoV-2 and other beta- and alphacoronaviruses may arise.

**Significance:** Understanding mechanisms of variation in coronaviruses is vital to control of their associated diseases. Beyond point mutations, insertions, deletions and recombination, we here describe for the first time intragenomic rearrangements and their relevance to changes in transmissibility, immune escape and/or virulence documented during the SARS-CoV-2 pandemic.

## Introduction

Coronaviruses (CoVs) are positive, singe stranded RNA viruses of the order Nidovirales, family Coronaviridae, subfamily Orthocoronavirinae, with four genera, namely alpha [α], beta [β], gamma [γ] and delta [δ], which have been further subdivided into 25 subgenera, including five for β-CoVs: Sarbeco-, Merbeco-, Embeco-, Nobeco- and Hibecovirus (Weiss and Navas-Martin 2005), and fifteen for α-CoVs: Luchaco-, Decaco-, Nyctacovirus, -Minunaco-, Pedaco-, Colaco-, Myotaco-, Duvinaco-Setraco-, Rhinaco-, Tegaco-, Minaco-, Sunaco-, Soracoviurs, and Amalacovirus (ICTV Coronaviridae Study Group. 2021). Seven CoVs infect humans; two of the α-genus (the Duvinacovirus hCoVs 229E & the Setracovirus NL63) and five of the β-genus: the Sarbecoviruses severe acute respiratory syndrome (SARS)-CoVs 1 and 2, the latter responsible for a pandemic since 2019 (Pollett et al. 2021; Jackson et al. 2021; VanInsberghe et al. 2021; Turkahia et al. 2021); the Merbecovirus Middle East respiratory syndrome (MERS) CoV; and the Embecoviruses hCoV-OC43 and -HKU1. Human CoVs have a zoonotic origin, with bats as key reservoir (Menachery et al. 2015) and possibly intermediate hosts (Fan et al. 2019; Pickering et al. 2021; Reusken et al. 2013; Song et al. 2005). Bat β-CoVs related to human CoVs belong to the Sarbeco-, Nobeco-, and Hibecovirus subgenera (Latinne et al. 2020; Wong et al. 2019; Woo et al. 2007).

Coronaviruses display substantial genomic plasticity and resilience (Amoutzias et al. 2022; Andersen et al. 2020) via recombination, point mutations, deletions, and insertions, which are reported to drive variant emergence, host range, gene expression, transmissibility, immune escape, and virulence (Decaro et al. 2009; Goldstein et al. 2021; Gussow et al. 2020; Simon-Loriere et al. 2011; Throne et al. 2022). The use of an RNA-dependent-RNA polymerase (RdRp)-driven template switching mechanism for transcription and control of structural and accessory gene expression in CoVs (Sawicki et al. 2007) has been reported to account for the high frequency of recombination (Amoutzias et al. 2022; Bobay et al. 2020; Boni et al. 2020; Forni et al. 2017, 2020; Lau et al. 2018; Makino et al. 1986; Simon-Loriere et al. 2011; Su et al. 2016; Yang et al. 2021).

In template switching, a leader transcription regulatory sequence (TRS-L; ACGAAC core in β-CoVs) (Wang et al. 2021) in the 5’-untranslated region (UTR) interacts with homologous TRS-body (B) elements upstream of viral genes in the last third of the genome (illustrated for SARS-CoV-2 in Figure 1A) (Bentley et al. 2013; Sawicki et al. 2007; Sola et al. 2015; Van Marle et al. 1995). Template switching renders the neighborhood of TRS-Bs, especially that for the spike gene, a recombination hotspot during viral transcription (Bobay et al. 2020; Boni et al. 2020; Forni et al. 2020; Graham et al. 2010, 2018; Goldstein et al. 2021; Lytras et al. 2022; Nikolaidis et al. 2021; Pollett et al. 2021; Yang et al. 2021).

**Figure 1.**
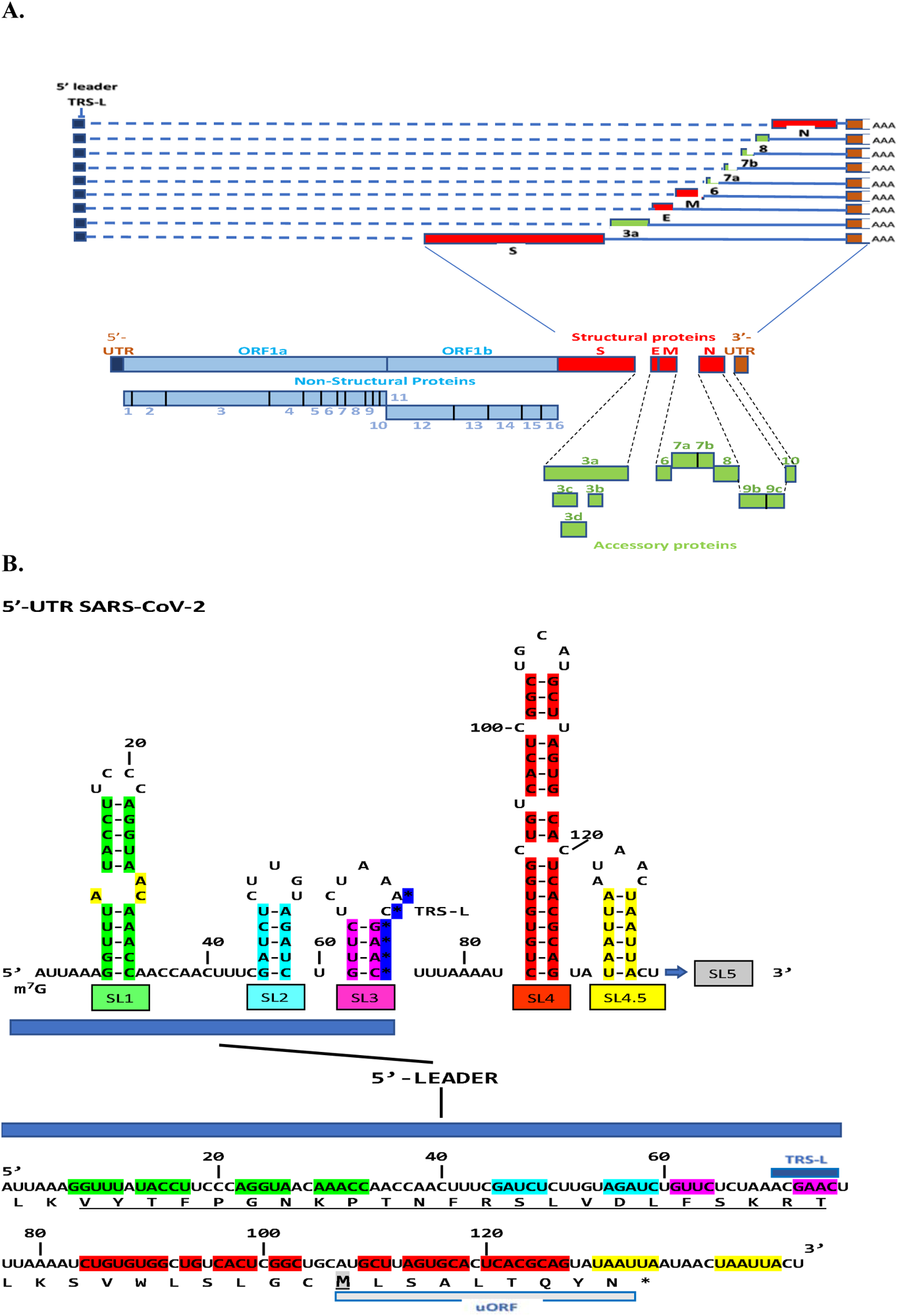
**A. Discontinuous synthesis of SARS-CoV-2 negative strand subgenomic RNA.** For the synthesis of subgenomic RNA, the leader transcription regulatory sequence (TRS-L, blue box) within the 5’-leader sequence interacts with homologous TRSs in the body (TRS-B) of the genome that precede structural (red boxes) and accessory (green boxes) genes. Overlapping genes for ORF3a (namely, ORF3b-d) and N (namely, ORF9b, c) that would be translated from the sgRNAs shown for ORF3a and N, respectively, are shown at the bottom. **B. Secondary structure of the 5’-UTR including the 5’-leader sequence and translated sequence of the 5’-leader sequence and beyond until the stop codon before stem-loop (SL)5.** The secondary structure of the 5’-UTR is shown as presented in Miao et al. (2021). The 5’ leader sequence extends from the cap structure (m^7^G) to the TRS-L and encompasses SL1-3, which have been associated with viral replication and gene expression. An open reading frame spans most of the 5’-UTR (Wuhan reference strain, NC_045512 shown) except for SL5, where ORF1ab starts. The segment of the open reading frame that is translocated wholly or partially in SARS-CoV-2 variants is underlined. At the nucleotide level, the segment includes the TRS-L, and at the amino acid level, the translated 5’-leader sequence and beyond includes a predicted upstream open reading frame (uORF, grey box) which has not been shown to be functional and whose initiation methionine [M] is shown in gray.

Viral subgenomic messenger RNAs contain a 5’-leader sequence that spans from the terminal 5’-cap (m^7^G) structure to the TRS-L and harbors three conserved stem-loop (SL1-3) regulatory elements of gene expression and replication (Figure 1B) (Madhugiri et al. 2018; Miao et al. 2021; Zhang et al. 1994). The TRS-L core sequence and the secondary structure of the leader sequence are conserved within but not among coronavirus genera (Rfam database: http://rfam.xfam.org/covid-19).

The entire 5’-leader nucleotide sequence of SARS-CoV-2, and beyond up to almost SL5 can be translated into a peptide sequence (Figure 1B), and although there is no evidence for the functionality of any open reading frame within the UTRs (Chen et al., 2010; Miao et al., 2021), the 5’-leader sequence is translated after most of it (nucleotides 8-80, including SL1-3 and TRS-L) is duplicated and translocated to the distal end of the accessory ORF6 gene of a SARS-CoV-2 variant with deleted ORFs 7a, 7b and 8 isolated from 3 patients in Hong Kong (Tse et al. 2021). We (Patarca and Haseltine 2021) also reported that a shorter portion of the 5’-leader sequence (nucleotides 50-75) is duplicated and translocated to the end of the accessory ORF8 gene of a USA variant generating a modified ORF-8 protein.

In the present study, using 5’-leader nucleotide sequences and amino acid sequences translated in the three reading frames as queries to search public databases, we document the presence of intragenomic rearrangements involving segments of the 5’-leader sequence in geographically and temporally diverse isolates of SARS-CoV-2. The intragenomic rearrangements modify the carboxyl-termini of ORF-8 (also in *Rhinolophus* bat Sarbecovirus β-CoVs) and ORF7b; the serine-arginine-rich region of the nucleocapsid protein, generating the well characterized R203K/G204R paired mutation; and two sites of the NiRAN domain of the RdRp (nsp12).

Beyond SARS-CoV-2, we found similar rearrangements of 5’-UTR leader sequence segments including the TRS-L in all subgenera of β-CoVs except for Hibecovirus (possibly secondary to the availability of only 3 sequences in GenBank). These rearrangements are in the intergenic region between ORFs 3 and 4a, and at the carboxyl-terminus of ORF4b of the Merbecovirus MERS-CoV; intergenic regions in the Embecoviruses hCoV-OC43 (between S and Ns5) and hCoV-HKU-1 (between S and NS4); and in the Y1 cytoplasmic tail domain of nsp3 of Nobecoviruses of African *Rousettus* and *Eidolon* bats. We also found intragenomic rearrangements in α-CoVs in nsp2 (Luchacovirus subgenus), nucleocapsid (Nyctacovirus subgenus), and ORF5b or ORF4b (Decacovirus subgenus). No rearrangements involving 5’-UTR sequences were detected for the β-CoV SARS-CoV-1; the other 12 subgenera of α-CoVs including hCoV-229E and hCoV-NL63 infecting humans; or δ (Andeco-, Buldeco-, and Herdecovirus subgenera) and γ CoVs (Brangaco-, Cegaco-, and Igacovirus subgenera) for which wild birds are the main reservoir (Wille and Holmes 2020; Woo et al. 2009).

The present study highlights an intragenomic source of variation involving duplication, inversion (in two α-CoVs subgenera) and translocation of 5’-UTR sequences to the body of the genome with implications on gene expression and immune escape of α- and β-CoVs in humans and bats causing mild-to moderate or severe disease in endemic, epidemic, and pandemic settings. Genome-wide annotations had revealed 1,516 nucleotide-level variations at different positions throughout the entire SARS-CoV-2 genome (Islam et al. 2020) and a recent study documented outspread variations of each of the six accessory proteins across six continents of all complete SARS-CoV-2 proteomes which was suggested to reflect effects on SARS-CoV-2 pathogenicity (Hassan et al. 2022). The intragenomic rearrangements involving 5’-UTR sequences described here, which in several cases affect highly conserved genes with a low propensity for recombination, may underlie the generation of variants homotypic with those of concern or interest and with differing pathogenic profiles.

## Results

Using the approaches described in the Materials and Methods section, we conducted a systematic analysis of SARS-CoV-2 and other coronaviruses and detected insertions involving 5’-UTR sequences at various locations in β- and α-CoVs, as described below by subgenus.

### Intragenomic rearrangements alter the carboxyl termini of ORF8 and ORF7b (Sarbecoviruses)

We had reported on a U.S. isolate of SARS-CoV-2 in which a segment encompassing nucleotides 50-75 of the 5’-UTR was duplicated and translocated to the end of the accessory ORF8 gene giving rise to an ORF8 protein with modified carboxyl-terminus encoded by the translocated 5’-UTR sequences (Patarca and Haseltine 2021). Figure 2 summarizes the results of our systematic search which revealed 240 similar insertions of various lengths of the same 5’-UTR sequence at various points in a stretch of 7 amino acids (_115_RVVLDFI_121_) of the ORF8 carboxyl-terminal sequence. As depicted in Supplemental legend to Figure 2, these internal rearrangements were detected in temporally and geographically diverse isolates, collected from March 2020 to December 2021 in 38 USA states, Bahrain, China, Kenya, and Pakistan, which is not exhaustive of what exists. All translocated 5’-UTR nucleotide sequence segments include TRS-L with variable extents of SL3 and SL2, that could affect expression of the nucleocapsid gene located immediately after the ORF8 gene (Thorne et al. 2022), and all insertions alter the carboxyl-terminus of ORF8. The analysis also revealed that the insertions in some isolates had further changes involving point mutations, deletions, and insertions. Moreover, as shown in Figure 3A, a similar 5’-UTR-derived insertion at the carboxyl-terminus of ORF8 is seen in five Sarbecovirus β-CoVs from what is considered the animal reservoir for SARS-CoV-2, the *Rhinolophus* (horseshoe) bats residing in Indochina and Southwest China (Temmam et al. 2021) all the way to England (Crook et al. 2021).

**Figure 2.**
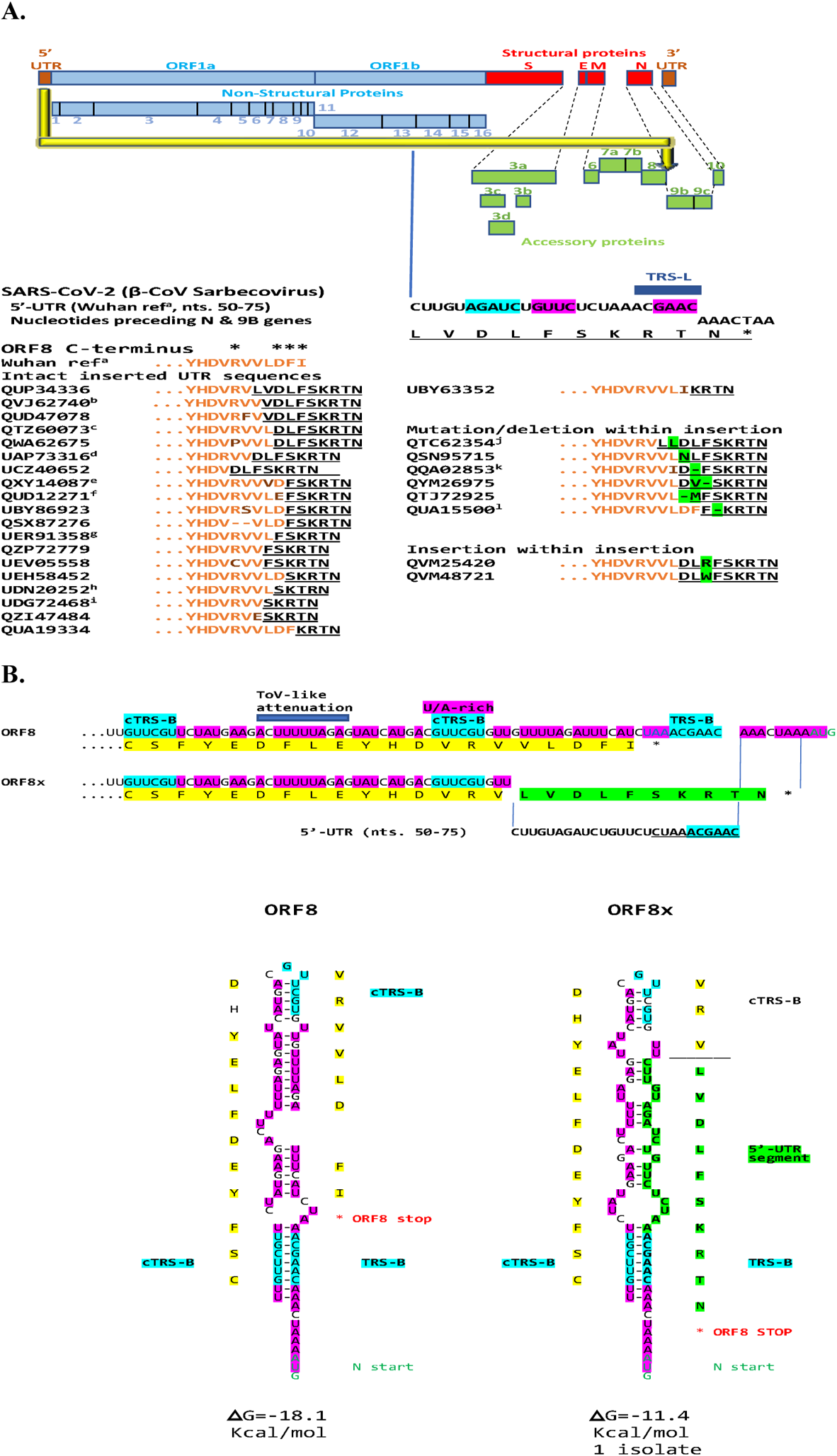
Modified carboxyl-termini of ORF8 encoded by a 5’-UTR-derived insertion in SARS-CoV-2. **A.** The largest 5’-UTR segment that was duplicated and translocated as an insertion to the carboxyl terminus of ORF8 is shown at the nucleotide and amino acid levels (latter underlined). All translocated 5’-UTR nucleotide sequence segments include TRS-L (dark blue box) with variable extents of SL3 (blue) and SL2 (red). Examples are shown, and corresponding similar sequences in GenBank as of February 20, 2022, are listed in the Supplemental legend to Figure 2. The C-terminus of ORF8 in the Wuhan reference strain is depicted using orange letters with mutations in ochre; the asterisks over the C-terminus sequence designate residues contributing to the covalent dimer interface (Arg115, Asp119, Phe120, Iso121; Flower et al. 2021). The 5’-UTR insertions are shown as underlined letters in black with mutations, deletions, and insertions within them highlighted in green. **B. Secondary structures of ORF8 RNA in reference strain and in that with longest 5’-UTR-derived intragenomic rearrangement.** Nucleotide and amino acid sequences of the carboxyl termini of ORF8 from Wuhan reference (NC_0445512) and from isolate QUP34336 (USA/Minnesota, 2021-04-05) with the longest 5’-UTR-derived sequence. ORF8 protein from the latter has modified carboxy terminus and is therefore designated ORF8x. Amino acid sequence from the reference strain is highlighted in yellow while that encoded by the duplicated and translocated 5’-UTR segment is highlighted in green. Stop codon of ORF8 is depicted with a red asterisk, and initiation codon of N is in green letters. TRS-B core sequence and complementary TRS-Bs in ORF8 and in ORF8x are highlighted in blue; and the uridine/adenosine tracks, including the torovirus-like attenuation sequence (Ujike and Taguchi. 2021) are highlighted in fuchsia. 5’-UTR nucleotide and amino acid sequences in ORF8X are in bold letters and highlighted in green.

**Figure 3.**
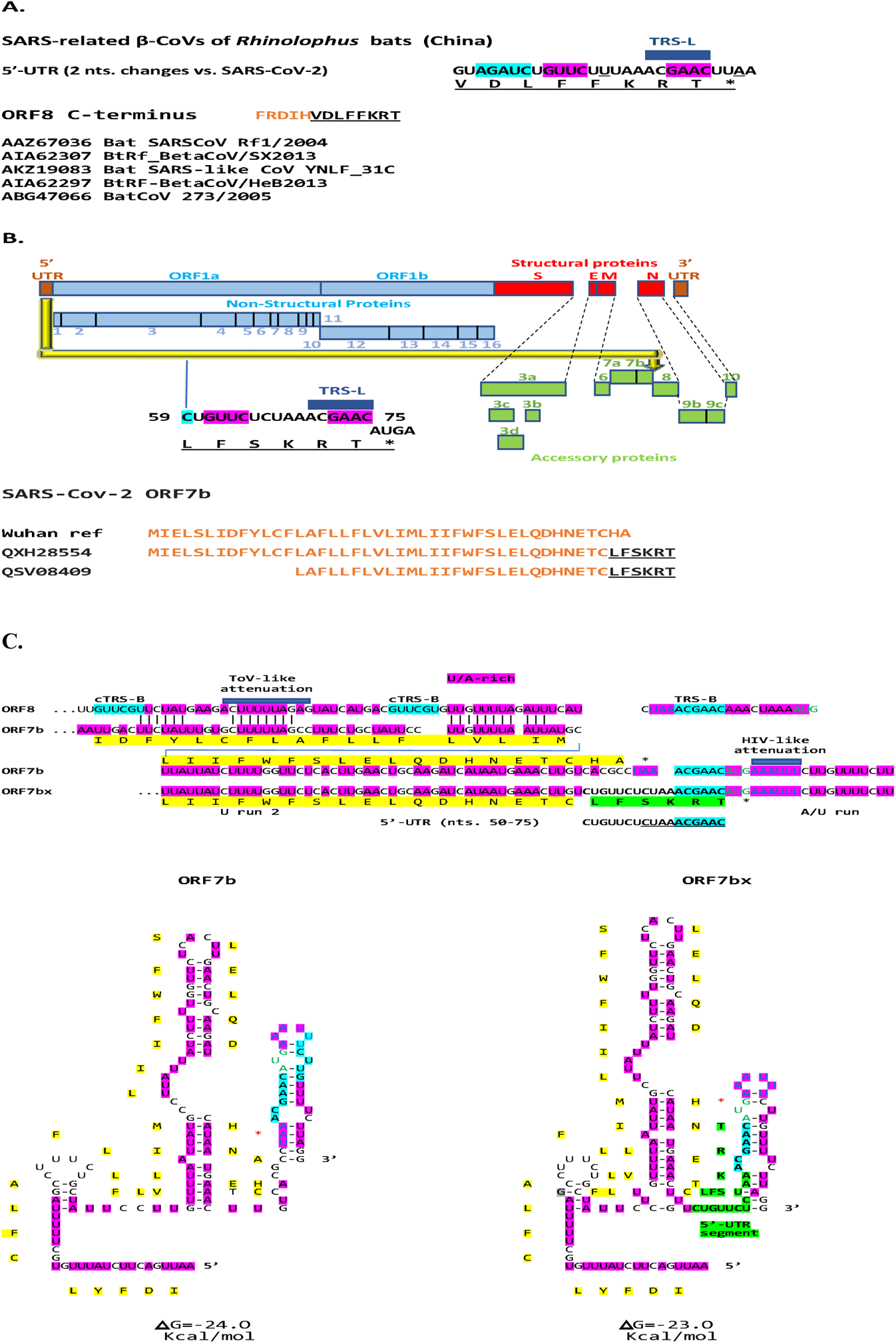
**A. Modified carboxyl-termini of ORF8 encoded by an insertion of a 5’-UTR segment in SARS-related-coronaviruses of *Rhinolophus* bats from China.** For SARS-related bat β-CoVs (BatSARSCoV Rf1/2004 and Bat CoV 273/2005 are subgroup 2b; Menachery et al. 2015), all inserted terminal sequences were the same. The nucleotide sequence of the inserted 5’-UTR segment differed from that of SARS-CoV-2 by two nucleotides: a C to U change (underlined) which translates into an amino acid change (serine [S] to phenylalanine [F]), and a U to A (underlined) which introduces a stop codon. **B. Modified carboxyl termini of ORF7b encoded by an insertion of a 5’-UTR segment in SARS-CoV-2.** The two isolates with modified ORF-7 proteins are QXH28554 (USA/Alabama, 2021/04/14), and QSV08409 (USA/California; 2021/02/26); the latter has a truncated ORF7b and the former a truncated ORF8. Color codes and abbreviations are as in Figures 1 and 2. **C. Secondary structures of RNAs for ORF7b and ORF7bx.** Color scheme is as in Figure 2B. An HIV-like attenuation sequence (Harrison et al. 1998) is also highlighted.

Figure 2C depicts the predicted secondary structure of ORF8 without and with (ORF8x) the 5’-UTR derived insertion. Both structures have similar predicted free minimum energy. The insertion involves the TRS-B sequence located in the intergenic region between ORF8 and N and is preceded by a uridine-adenosine (U/A)-rich region including a sequence similar to that of the torovirus attenuation sequence (Ujike and Taguchi 2021), which like TRS-B, could cause the RNA-dependent RNA polymerase to pause, thereby facilitating the intragenomic rearrangement as it is theorized to do during subgenomic RNA synthesis. Supplemental Figure 1 shows the predicted RNA structures of the most common ORF8x variants with similar predicted minimum free energy, and Supplemental Figure 2 shows an alternative RNA structure involving the interaction between the TRS-B in the intergenic region with a second and closer complementary TRS-B, yielding a similar predicted minimum free energy.

Crystal structure of ORF8 of SARS-CoV-2 had revealed a ∼60-residue core like that of SARS-CoV-2 ORF7a (from which ORF8 has been postulated to originate by non-homologous recombination) (Neches et al. 2021) with the addition of two dimerization interfaces, one covalent and the other noncovalent, unique to SARS-CoV-2 ORF8 (Flores et al. 2021). In the C-terminus of ORF8 that is altered by 5’-UTR-derived insertions (i.e., _115_RVVLDFI_121_), R115, D119, F120, and I121 contribute to the covalent dimer interface (marked with asterisks in Figure 2) with R115 and D119 forming salt bridges that flank a central hydrophobic core in which V117 interacts with its symmetry-related counterpart (Flores et al. 2021).

How the C-terminal insertions and changes therein affect the dimerization of ORF8 remains to be determined and described functions for ORF8 remain a matter of debate (Redondo et al. 2021). However, the changes caused by insertions may contribute to immune evasion by SARS-CoV-2 by affecting the interactions of ORF8 as a glycoprotein homodimer with intracellular transport signaling, leading to down-regulation of MHC-I by selective targeting for lysosomal degradation via autophagy (Zhang et al. 2021), and/or extracellular signaling (Matsuoka et al. 2022) involving interferon-I signaling (Li et al. 2020), mitogen-activated protein kinases growth pathways (Valcarcel et al. 2021), the tumor growth factor-β1 signaling cascade (Stukalov et al. 2021) and interleukin-17 signaling promoting inflammation and contributing to the COVID-19-associated cytokine storm (Lin et al. 2021).

The carboxyl-terminal region may include T- and/or B-cell epitopes that may be affected by the variations described. To this end, approximately 5% of CD4+ T cells in most COVID-19 cases are specific for ORF8, and ORF8 accounts for 10% of CD8+ T cell reactivity in COVID-19 recovered subjects (Gordon et al. 2020; Griffoni et al. 2020). Another possible effect of the insertions stems from the fact that anti-ORF8 antibodies are detected in both symptomatic and asymptomatic patients early during infection by SARS-CoV-2 (Hachim et al. 2020; Wang et al. 2020) and diagnostic assays for SARS-CoV-2 infection that target only accessory genes or proteins such as ORF8 may be affected (Tse et al. 2021).

A shorter segment of the SARS-CoV-2 5’-UTR leader sequence (nts. 57 to 95, including TRS-L and SL3) than that described for ORF8 insertions was also duplicated and translocated to the end of ORF7b in two SARS-CoV-2 isolates (Figure 3B), one with a truncated ORF7b and the other with a truncated ORF8, which may have favored the internal rearrangements. Figure 3C shows the predicted secondary structures of the region with the intragenomic rearrangement, and as that of ORF8, involves the intergenic TRS-B sequence which is preceded and followed by a U/A-rich region, in this case also incorporating an HIV-like attenuation sequence (AAAUUU; Harrison et al. 1998). Figure 3C also shows a region of similarity between ORF8 and ORF7b which precedes the intragenomic region. The function of the SARS-CoV-2 ORF7b remains to be determined and has been suggested to mediate tumor necrosis factor-α-induced apoptosis based on cell culture data (Yang R et al. 2021) and theoretically the dysfunction of olfactory receptors by triggering autoimmunity (Khavison et al. 2020).

### Intragenomic rearrangements alter the serine-arginine-rich region of the N protein (SARS-CoV-2)

In terms of structural proteins of SARS-CoV-2, we found a similar segment of the 5’-UTR corresponding to the leader sequence (nucleotides 56 to 76 of the Wuhan reference strain [NC_045512], including TRS-L, SL3 and part of SL2, and encoding the 7-amino acid sequence DLFSKRT) within the N gene at the end of its SR region, as exemplified by isolate QTO33828 (USA/Texas, Figure 4A). The 5’-UTR-derived segment changes 5 of 7 positions, including R203K/G204R, which are known to be frequent co-occurring mutations in the N protein; however, the rest of the N protein sequences are well conserved with only 1 or 2 amino acid differences in the isolates identified. In another set of SARS-CoV-2 isolates, as exemplified by isolate EPI-ISL_3434731 (Brazil/Espirito Santo) in Figure 4A, the same 5’-UTR-derived sequence is present in N but without the leucine (L) residue and the phenylalanine (F) changed to serine (S), more closely approaching the Wuhan reference strain sequence.

**Figure 4.**
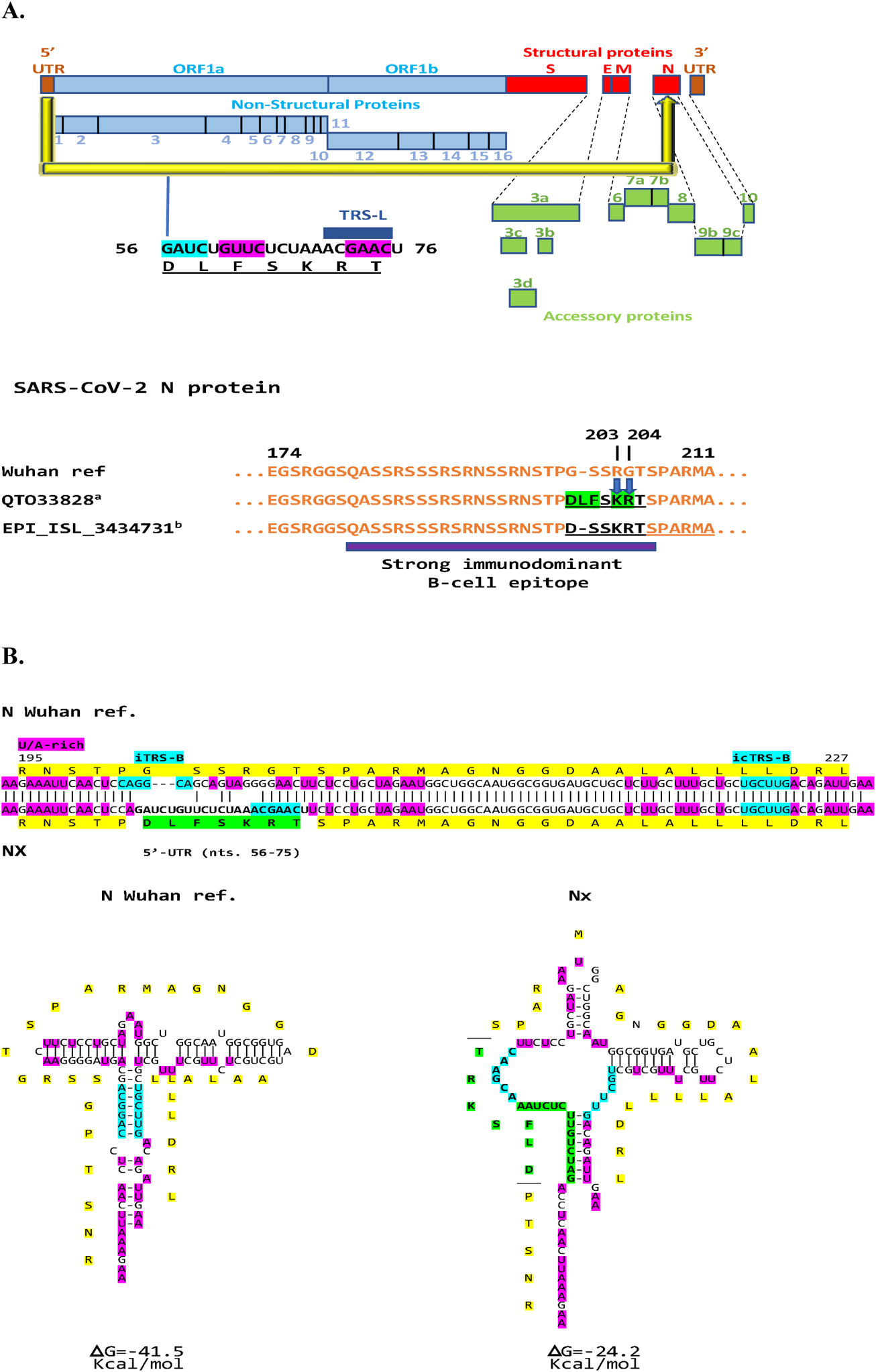
**A. Insertion of 5’-UTR segment into the serine-rich region of the nucleocapsid (N) in SARS-CoV-2.** The R203K and G204R amino acid substitutions (blue arrows) which are commonly present concomitantly are encoded in this case by the insertion of a 5’-TR segment into the SR-rich region of the N protein at the end of a strong immunodominant B-cell epitope (purple box; Oliveira et al. 2020). B. **Secondary structures of RNAs for N and Nx.** Color scheme is as in preceding Figures.

The predicted RNA structures of the N protein without and with (Nx) the 5’-UTR-derived insertion are shown in Figure 4C, with that for Nx being more stable with almost half the minimum free energy. Although there is no TRS-B in the N region where the intragenomic rearrangement was found, there is an inverted TRS-B that can pair with a complementary inverted TRS-B, both surrounded by U/A-rich regions which could facilitate the intragenomic rearrangement.

In total, 37 SARS-CoV-2 isolates had 5’-UTR-derived sequences in their N gene, in contrast to ∼336,000 isolates with either R203K or G204K as per NCBI Virus (mutations in SARS-CoV-2 SRA data); most were isolates of the variant of concern gamma GR/501Yv3 (P1) lineage (first detected in Brazil and Japan) from Brazil, Chile, and Peru, but also alpha (B.1.1.7; first detected in Great Britain) from USA and Canada (Supplemental legend to Figure 4). The R203K/G204R co-mutation has been associated with B.1.1.7 (alpha) lineage emergence, which along with variants with the co-mutation including the P1 (gamma) lineage (Franco-Muñoz et al. 2020), possess a replication advantage over the preceding lineages and show increased nucleocapsid phosphorylation, infectivity, replication, virulence, fitness, and pathogenesis as documented in a hamster model, human cells, and COVID-19 patients including an analysis of association between COVID-19 severity and sample frequency of R203K/G204R co-mutations (Johnson et al. 2021; Mourier et al. 2022; Wu et al. 2021).

The nucleocapsid is the most abundant protein in CoVs, interacts with membrane protein (He et al 2004; Lu et al. 2021), self-associates to provide for efficient viral assembly (Yao et al. 2020), binds viral RNA (McBride et al. 2014) and has been involved in circularization of the murine hepatitis virus genome via interaction with 3’- and 5’-UTR sequences which may facilitate template switching during subgenomic RNA synthesis (Lo et al. 2019). Phosphorylation transforms N-viral RNA condensates into liquid-like droplets, which may provide a cytoplasmic-like compartment to support the protein’s function in viral genome replication (Carlson et al. 2020; Lu et al. 2021).

The phosphorylation-rich stretch encompassing amino acid residues 180 to 210 (SR region) in which the 5’-UTR-derived sequences were found, serves as a key regulatory hub in N protein function within a central disordered linker for dimerization and oligomerization of the N protein, which is phosphorylated early in infection at multiple sites by cytoplasmic kinases (reviewed in Carlson et al. 2020). Serine 202 (numbering of reference Wuhan strain), which is phosphorylated by GSK-3, is conserved in the 5’-UTR-derived sequence next to the R203K/G204R co-mutation, as is threonine 205, which is phosphorylated by PKA (Kemp et al. 1977; Kennelly et al. 1991). R203 and G204 mutations affect the phosphorylation of serines 202 and 206 in turn affecting binding to protein 14-3-3 and replication, transcription, and packaging of the SARS-CoV-2 genome (Surjit et al. 2005; Tugaeva et al. 2021; Tung and Limtung 2021).

The N gene displays rapid and high expression, high sequence conservation, and a low propensity for recombination (Dutta et al. 2020; Jaroszewski et al. 2021; Nikolaidis et al. 2021). However, it can show variation driven by internal rearrangement which does not affect the length of the protein. The N protein is highly immunogenic, and its amino acid sequence is largely conserved, with the SR region being a strong immunodominant B-cell epitope (Oliveira et al. 2020) as highlighted in Figure 4A.

### Intragenomic rearrangements alter the Nidovirus RNA-dependent RNA polymerase associated nucleotidyl transferase (NiRAN) domain (SARS-CoV-2)

Another example of intragenomic rearrangement is the presence of the translated sequence (DLFSK) of a shorter segment of 5’ UTR (nts. 56-70 in Wuhan reference strain, including parts of SL2 and SL3 but not TRS-L) at amino acids 36-40 of the NiRAN domain of the viral RdRp (nsp12) in isolates QVL75820 (EPI_ISL_1209225, USA/Seattle, 2021-03-28; lineage: B.1.2 [Pango v.3.1.20 2022-02-02]) and EPI_ISL_1524008 (USA/Washington, 2021-03-28; VOC Alpha GRY (B.1.1.7+Q.*) first detected in the UK) and at amino acids 146-150 in isolates UFT72204 (EPI_ISL_6912949, USA/Colorado, 2021-10-27; VOC Delta GK [B.1.617.2+AY.*] first detected in India), EPI_ISL_1384819 (India/Maharashtra, 2021-02-12; lineage: B.1.540 [Pango v.3.1.20 2022-02-02]) and EPI_ISL_1703925 (India/Maharashtra, 2021-02-07; B.1.540 lineage), respectively (Figure 5A). The latter strains have only one amino acid change outside of the insertions relative to the Wuhan reference strain. A subsegment of 5’-UTR (nts. 62-70) translated as FSK is present at the more proximal site (amino acids 38-40) in 230 isolates isolated from diverse populations at various times (listed in Supplemental legend to Figure 5) and exemplified by isolate UHP90975 [USA/Wisconsin, 2021-12-13] in Figure 5A. Isolate QZM71485 (USA/New York, 2021-08-05) exemplifies isolates with the FSK sequence at the more distal site (amino acids 148-150). Examples of the most common single amino acid changes in overlapping segments of other isolates are listed as comparators, and they have similar or lower frequency than those of the 5’-UTR-derived segments. However, the Wuhan reference strain sequence corresponding to the areas with 5’-UTR sequences is the most abundant among SARS-CoV-2 isolates.

**Figure 5.**
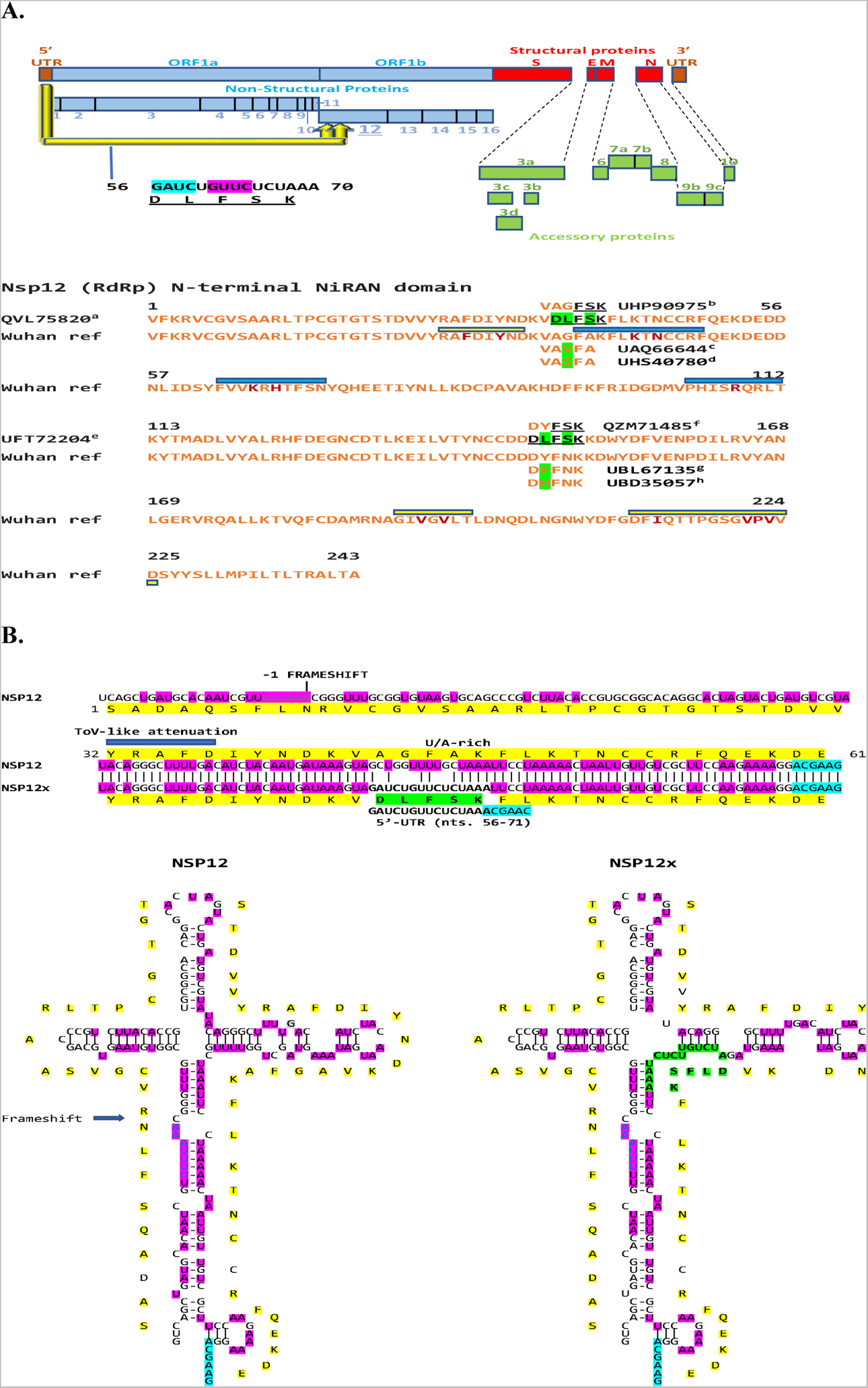
**A. Insertions of a 5’-UTR sequence into two sites within the Nidovirus RdRp associated nucleotidyl transferase (NiRAN) domain of the RNA-dependent RNA polymerase (nsp12) of SARS-CoV-2.** Examples of isolates with 5’-UTR-derived sequences at the proximal and distal sites are provided in the figure and a full listing is provided in the Supplemental legend to Figure 6, as is a listing of variants with single amino acid changes relative to the Wuhan reference strain in the segment corresponding to the insertion. The Wuhan reference strain sequence corresponding to the insertion areas is the most abundant among SARS-CoV-2 isolates. The nsp12-nsp9 interface regions are shown with yellow bars and key residues therein with ochre letters, while the contact regions with GDP are indicated with blue boxes and key residues therein in ochre. **B. Secondary structures for RNAs in the proximal sites in nsp12 and nsp12x.** Color scheme is as in previous figures. The site for -1 frameshifting is also highlighted.

Genes encoding components of the replication-transcription complex, such as the RdRp (nsp12) (Hartenian et al. 2020; Lauber et al. 2013), are highly conserved and have a low propensity for recombination among CoVs (Nikolaidis et al. 2021). The nsp12 NiRAN domain is one of the five replicative peptides that are common to all Nidovirales and used for species demarcation because it is not involved in cross-species homologous recombination (Gorbalenya et al. 2020). However, as in other examples here of conserved genes, it is involved in intragenomic rearrangements of 5’-UTR-derived sequences. Figure 5B shows the predicted structure RNA structures for the proximal site of intragenomic rearrangement in nsp12 and nsp12x (with 5’-UTR-derived sequence). As in the case of the example in the intragenomic rearrangement in N there is no TRS-B at the site of intragenomic rearrangement, which is however preceded by a torovirus-like attenuation sequence within a U/A-rich region which may facilitate pausing of the RdRp.

The NiRAN domain of nsp12 is involved in the NMPylation of nsp9 (Slanina et al. 2021) during the formation of the replication-transcription complex (interface regions [Yan et al. 2021] are shown with yellow bars and key residues therein with ochre letters in Figure 6). The 5’-UTR-derived sequence at the proximal site in the nsp12 NiRAN domain overlaps with one of the interface regions with nsp9 but does not affect key interface residues or alter the charge distribution of amino acid side chains in the overlap region. The nsp12 NiRAN domain also exhibits a kinase/phosphotransferase like activity (Dwivedy et al. 2021), is involved in protein-primed initiation of RNA synthesis (Lehmann et al. 2015) and catalyzes the formation of the cap core structure (GpppA; contact regions with GDP [Yan et al. 2021] indicated with blue boxes and key residues therein in ochre in Figure 5A) (Park et al. 2022). The 5’-UTR-derived sequence at the proximal site in nsp12 NiRAn domain is close to the first contact region with GDP.

**Figure 6.**
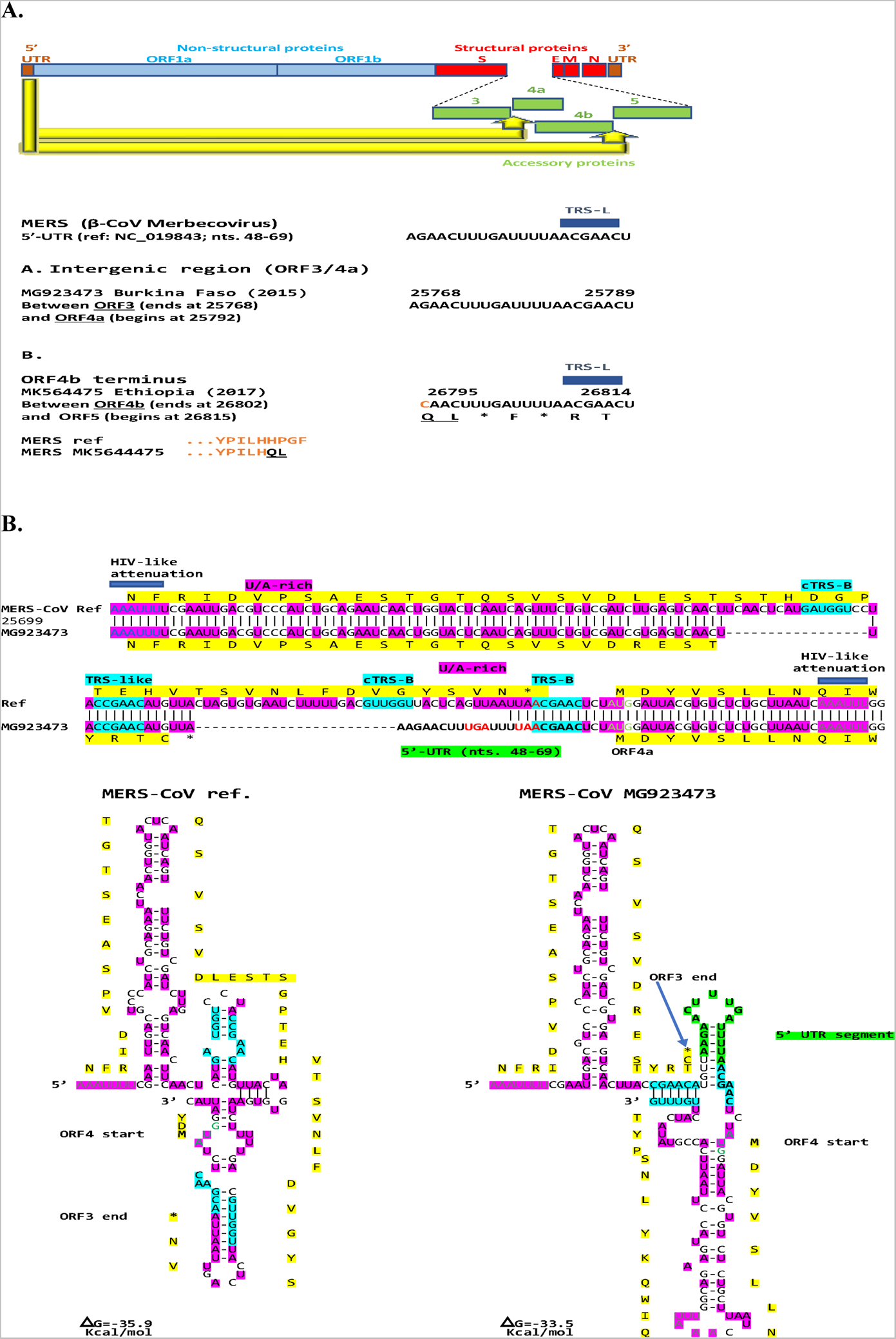
**A. Intragenomic rearrangement with 5’-UTR sequences present in the intergenic regions between p3 and 4a as well as between p4b and p5 of the Merbecovirus Middle East respiratory syndrome (MERS)-CoV.** A segment of the 5’-UTR of the MERS-CoV including TRS-L and part of the second of the two stem-loops is present in the intergenic region between p3 and p4a in isolate MG923473 (Burkina Faso, 2015) and between p4b and p5 affecting the carboxyl-terminal end of ORF4b in isolate MK564475 (Ethiopia, 2017). In the latter case, the last 4 amino acids (HPGF) of ORF4b in the reference MERS-CoV sequence (NC_019843) are replaced by two amino acids (QL). The Q residue is encoded by a cytosine present in the reference sequence (indicated in orange color) and two adenosines incorporated by the 5’-UTR-drived sequence. **B. RNA secondary structures of the intergenic region between p3 and p4a in MERS-CoV without and with intragenomic rearrangement.** Color scheme is as in previous figures.

### Intragenomic rearrangements in **β**-CoVs of Merbecovirus, Embecovirus, and Nobecovirus subgenera

#### Merbecovirus

As shown in Figure 6A, a segment of the 5’-UTR of the β-CoV Merbecovirus MERS-CoV including TRS-L and part of the second of the two stem-loops is present in the intergenic region between p3 and p4a in isolate MG923473 (Burkina Faso, 2015) and at the carboxyl-terminal end of p4b in isolate MK564475 (Ethiopia, 2017). In the latter case, the last 4 amino acids (HPGF) of p4b in the reference MERS-CoV sequence (NC_019843) are replaced by two amino acids (QL). The Q residue is encoded by a cytosine present in the reference sequence (indicated in orange in Fig. 6A) and two adenosines incorporated by the 5’-UTR-derived sequence. Figure 6B depicts the predicted RNA secondary structures without and with the insertion corresponding to the intragenomic rearrangement between p3 and p4b. The structures have similar predicted free minimum energy, and the rearrangement involves the intergenic TRS-B sequence which is preceded and succeeded by torovirus-like attenuation sequences.

p4a, a double stranded RNA-binding protein, as well as p4b and p5 of MERS-CoV are type-I IFN antagonists (Liu et al. 2014; Matthews et al. 2014; Niemeyer et al. 2913; Siu et al. 2014). p4a prevents dsRNA formed during viral replication from binding to the cellular dsRNA-binding protein PACT and activating the cellular dsRNA sensors RIG-I and MDA5 (Niemeyer et al. 2013; Siu et al. 2014). p4a is the strongest in counteracting the antiviral effects of IFN via inhibition of both its production and Interferon-Stimulated Response Element (ISRE) promoter element signaling pathways (Yang et al. 2013). Therefore, the intragenomic rearrangements found in MERS-CoV may facilitate immune evasion by bringing regulatory sequences to the intergenomic regions preceding the 4a and 5 genes and facilitating their expression.

#### Embecoviruses

The β-CoV Embecovirus hCoV-HKU1 is a sister taxon to murine hepatitis virus and rat sialodacyoadenitis virus (Corman et al. 2018). Out of 48 HKU-1 isolates in GenBank, a 5’-UTR sequence including TRS-L, SL3 and most of SL2 (nucleotides 42-74 in hCoV-HKU-1 references NC_006577 and AY597011) is present in 31 isolates (65%) between the S and Ns4 genes (Figure 7A). The hCoV-HKU-1 NS4 protein is structurally similar to the hCoV-OC43 ns5a protein whose function is detailed below.

**Figure 7.**
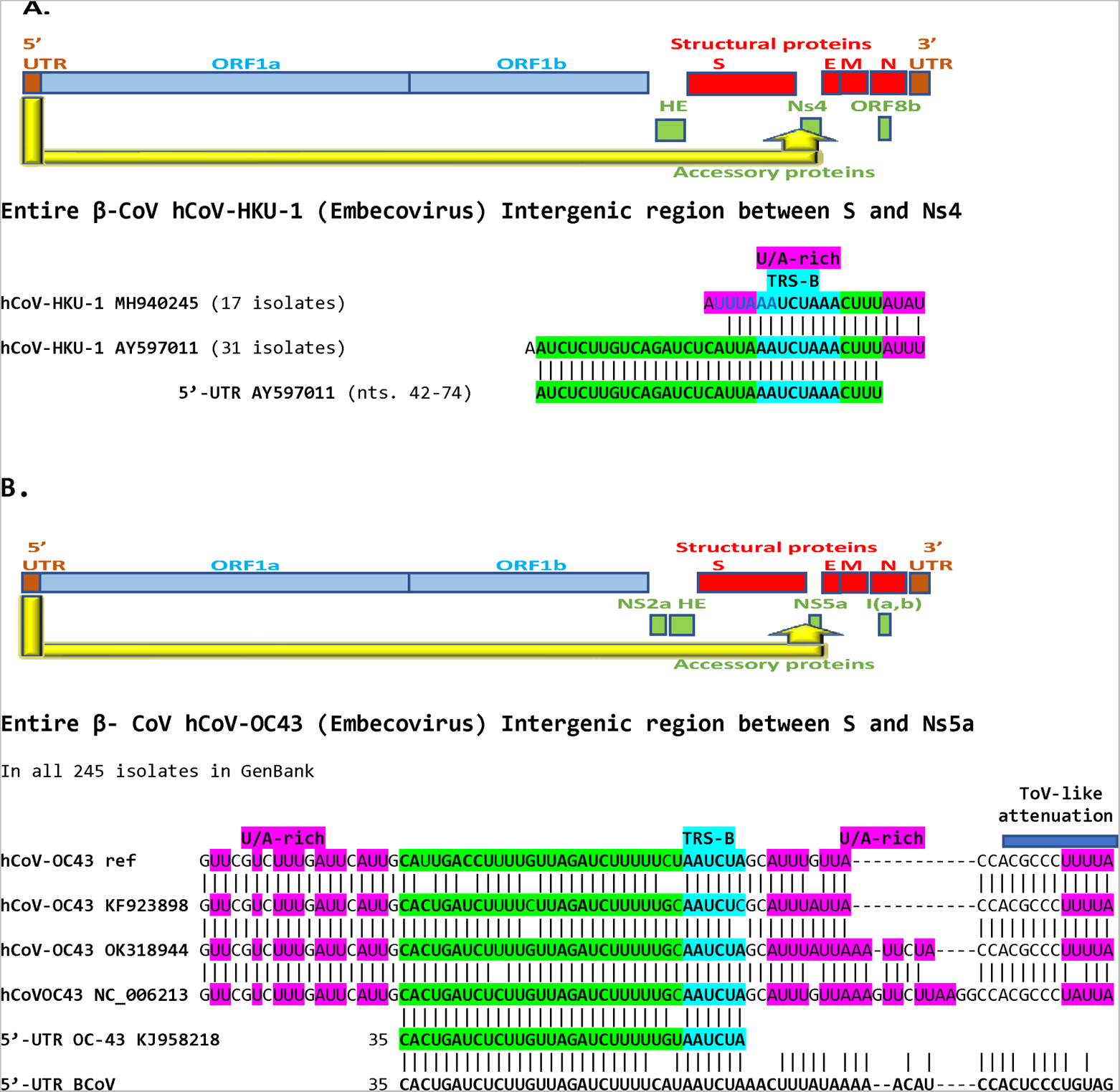
**A. Presence of hCoV-HKU-1 (β-CoV Embecovirus) of 5’-UTR-derived sequence in the intergenic region between the spike (S) and the Ns4 genes.** The hCoV-OC43 5’-UTR sequence inserted is identical to that of bovine coronavirus (BCoV) 5’-UTR (shown at the bottom of the figure) except for one nucleotide (an adenosine [A] instead of a guanosine [G] in BCoV). 31 out of 48 variants (65%) in GenBank have this intragenomic rearrangement. **B. Presence in in the intergenic region between the S and Ns5a genes hCoV-OC43 (β-CoV Embevovirus) of sequences of various lengths of the same 5’-UTR region.** As in the case of hCoV-HKU1, the 5’-UTR segment translocated to the intergenic region between S and Ns5a of hCoV-OC43 variants is similar to that of BCoV. Differences among them(all 245 isolates in GenBank) are distally to the TRS-B and involve various extents of sequences similar to the 5’-UTR of BCoV.

All 245 full genome isolates of the β-CoV Embevovirus hCoV-OC43 in GenBank had 5’-UTR-leader derived sequences (largest spanning nucleotides 35-67 of the hCoV-OC43 reference strain KJ958218) between the spike (S) and Nsp5a genes (Figure 7B). The insertions did not affect the protein sequences of either S or Nsp5a. The hCoV-OC43 5’-UTR sequence inserted is identical to that of bovine coronavirus (BCoV) 5’-UTR except for one nucleotide (an adenosine substituted by a guanosine in BCoV) up to the TRS-B, and sequences of varying length after the TRS-B show similarities to BCoV 5’-UTR, which is consistent with a most probable bovine or swine coronavirus origin for hCoV-OC43 (Vijgen et al. 2005). The 5’-UTR-derived insertion sequence is also present in a molecularly characterized cloned hCoV-OC43 S protein gene (Mounir et al. 1993).

hCoV-OC43 ns5a, as well as ns2a, M, or N protein significantly reduced the transcriptional activity of ISRE, IFN-β promoter, and NF-κB-RE following challenge of human embryonic kidney 293 (HEK-293) cells with Sendai virus, IFN-α or tumor necrosis factor-α (Beidas et al. 2018a). Like SARS-CoVs and MERS-CoV, hCoV-OC43 can downregulate the transcription of genes critical for the activation of different antiviral signaling pathways (Beidas et al., 2018b), and the intragenomic rearrangements described in the intergenic region preceding hCov-OC43 ns5a may facilitate immune evasion as was mentioned above for other immunomodulatory accessory proteins.

The Spike (S) gene encodes a structural protein that binds to the host receptors and determines cell tropism as well as the host range. As mentioned in the Introduction, the neighborhood of the spike gene, particularly the region before the S gene, is a hotspot for modular intertypic homologous and non-homologous recombination in coronavirus genomes (Nikolaidis 2021). In the cases described above for hCoV-HKU-1 and hCoV-OC43, intragenomic rearrangements involved the intergenic region at the end of the S gene highlighting a potential source of regulatory sequences that may affect expression of adjoining genes.

#### Nobecoviruses

An intragenomic rearrangement involving a 5’-UTR sequence (nucleotides 1-55) to the C-terminal cytoplasmic Y1 domain of nsp3 (nucleotides 6837-6891; amino acids 2188-2205), is seen in the β-CoV subgenus Nobecovirus of African bats, namely isolates MIZ240 (OK067321) and MIZ178 (OK067320) from *Rousettus madagascariensis* bats and isolates CMR900 (MG693169; protein: AWV67046), CMR705-P13 (MG693172, protein: AWV67070), and unclassified (NC_048212) from *Eidolon helvum* bats (Cameroon). Using the translated nucleotide sequence as query, the following additional isolates were detected: *Eidolon helvum* (Cameroon) isolates CMR704-P12 (YP_009824989 and YP_009824988), and CMR891-892 (AWV67062). The 5’-UTR sequence involved in this intragenomic rearrangement does not include the TRS-L and includes a stem-loop structure highlighted in grey in Figure 8A. The position of the translated sequence of the 5’-UTR identical sequence is amino acids 2188-2205, which corresponds to amino acids 1567-1584 in SARS-CoV-2 nsp3. Figure 8B depicts the predicted secondary structures of nsp3 and nsp3x with the intragenomic rearrangement. Both structures have similar predicted minimum free energy. Although there are no TRS-B sequences present in this region the rearrangement takes place adjacent to an inverted complementary TRS-B within a U/A-rich region.

**Figure 8.**
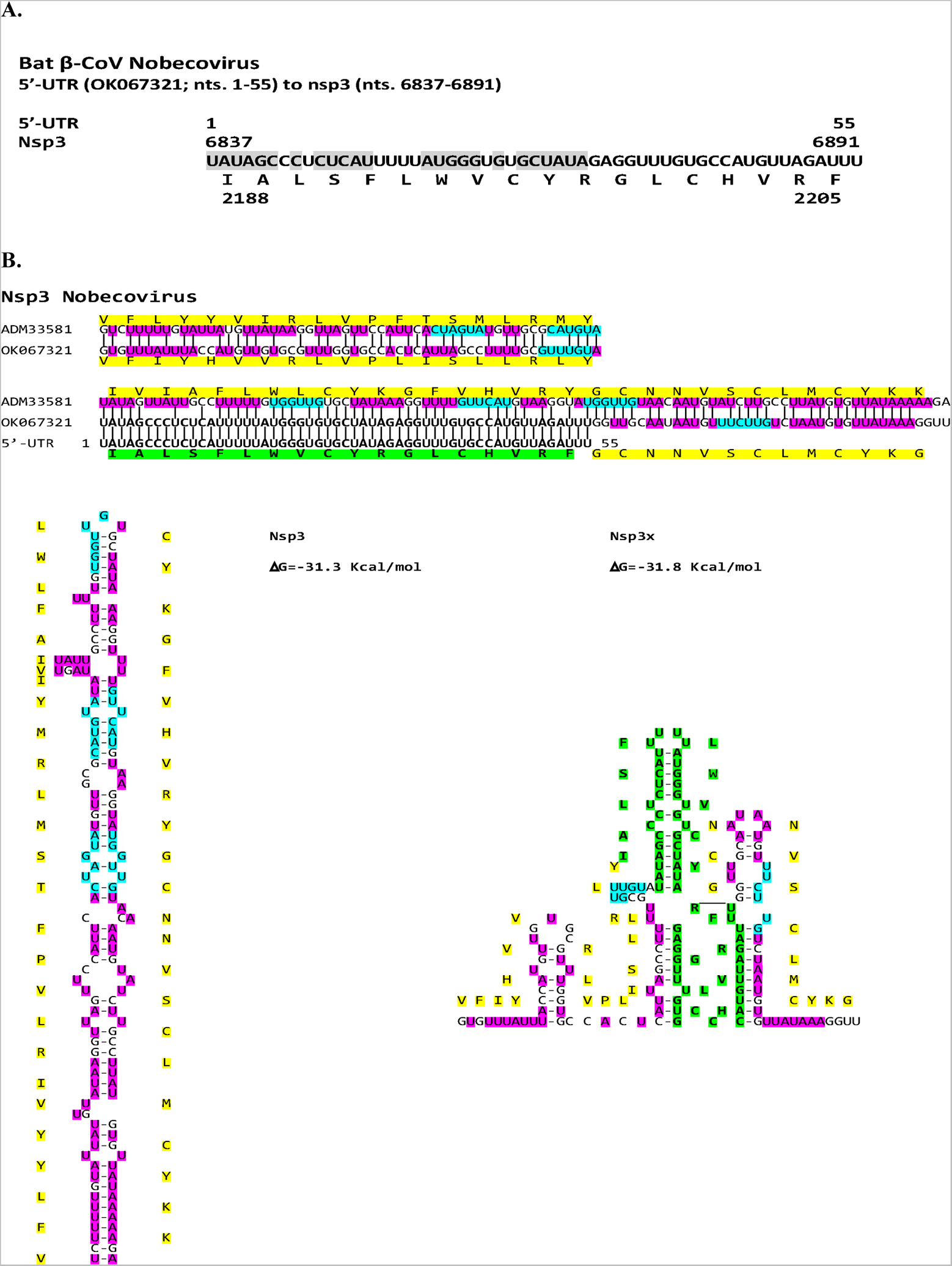
**A. Presence of 5’-UTR sequence in the Bat β-CoV Nobecovirus nsp3 gene.** All variants in GenBank with this intragenomic rearrangement are listed in the Supplemental legend to Figure 8. **B. Secondary structures of the RNAs of nsp3 and nsp3x.** Color scheme is as in previous figures.

Nsp3 protein, the largest protein encoded by CoVs encompasses up to 16 modular domains. The N-terminal cytosolic domains include a mono-ADP-ribosylhydrolase (Alhammad et al. 2021), a papain-like protease (Lei et al. 2018), and a scaffold region that participates in replication-transcription complex assembly (Imbert et al. 2008). After the latter domains, there are two transmembrane domains (TM1 and TM2) with an endoplasmic reticulum luminal loop (Ecto3) between them, and two cytosolic domains (Y1 and CoV-Y) following TM2. The nsp3 segment encoded by the 5’-UTR-derived sequence falls in the cytosolic domain Y1. Nsp3C anchors nsp3 to the endoplasmic reticulum membrane and induces membrane rearrangement leading to double membrane vesicle formation via a yet unknown molecular mechanism (Angelini et al. 2013; Hagemeijer et al. 2014). Although there are structural data on the CoV-Y domain (Pustovalova et al. 2021), its function is unknown as is that of the Y1 domain. Therefore, although the nsp3 sequence is well conserved among bat Nobecoviruses, the significance of the nsp3 segment encoded by the 5’-UTR-derived sequence, which might affect double vesicle membrane formation, remains to be determined.

### Intragenomic rearrangement in nsp2 of rodent α-CoVs subgenus Luchacovirus

As shown in Figure 9A, a segment of the 5’-UTR (nts. 1-119) of the Luchacovirus AcCoV-JC34 (KX964649; isolated in China, 2011-10 from the rodent *Apodemus chevieri*) was duplicated, inverted, and translocated to the genomic region encoding the nonstructural protein nsp2 (nts. 1679-1760). The latter sequence in nsp2 differs by only one nucleotide from that in the 5’-UTR (99% similarity), and it is also present with varying degrees of similarity in other rodents. Two examples are shown in Figure 9A for isolates from rat (Lucheng Rn rat CoV isolate Ruian 83; MT820626; isolated from *Rattus norvegicus* in China, 2014, 76% similar), and mouse (Fievel mouse CoV strain FiCoV/UMN2020 (OK655840; isolated from *Mus musculus* in USA, 2018, 59% similar). Other isolates (listed by rodent of origin) with intragenomic rearrangements in nsp2 with nucleotide sequences up to 75% similar to the 5’-UTR-derived sequences include: *Apodemus chevrieri* (MT820625, China, 2015, 93% similar); *Apodemus agrarium* (MZ328302, China, 2016, 93% similar); *Eothenomys miletus* (MT820627, China, 2014, 81% similar); *Eothynomys melanogaster* (KY370054, China, 2015-12, 79% similar); *Myodes rufocanus* (KY370045, China, 2014-08, 79% similar); *Rattus losea* (KY370050, China, 2015-05, 78% similar); *Rattus norvegicus* (MK163627, United Kingdom, 2014-06-23, 78% similar; NC_032730, China, 2013; MT549854, China, 2016-12, 76% similar; MW802582, China, 2017-03-07, 76% similar); and *Brylmys bowersi* (MZ328301, China, 2016, 77% similar).

**Figure 9.**
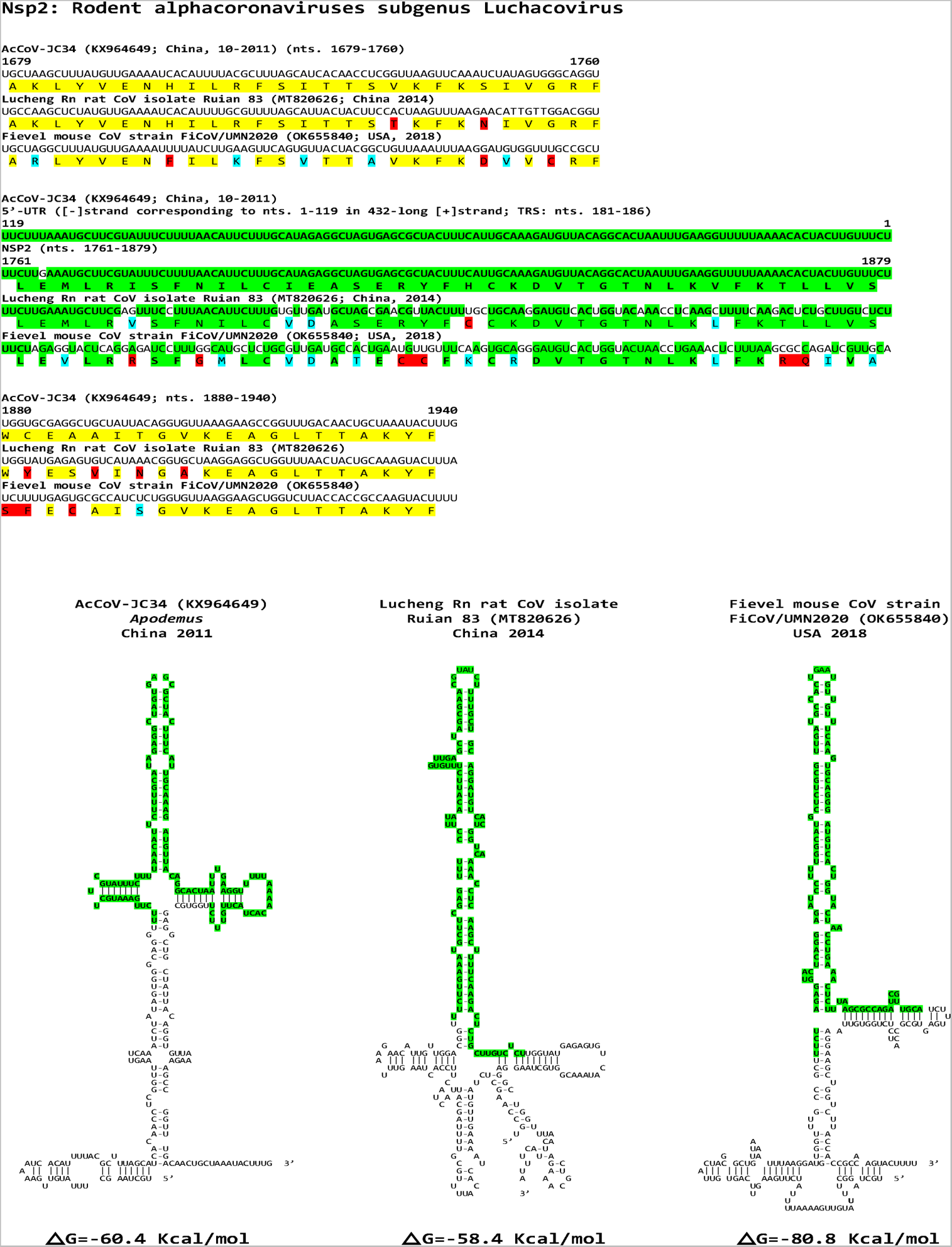
Intragenomic rearrangement in nsp2 of rodent alphacoronaviruses subgenus Luchacovirus. Nucleotide and amino acid sequences of intragenomic rearrangements are shown. Region derived from 5’-UTR (negative strand) is highlighted in green. Conservative amino acid substitutions are highlighted in blue while non-conservative ones are highlighted in red. Two examples of isolates (listed by rodent of origin) with nsp2 nucleotide sequences up to 75% similar to the 5’-UTR-derived sequences are shown. There appears to be a temporal gradient with the most similar sequence (99%) in isolate KX964649 (China, 2011-10) to the least similar (59%) in isolate OK655840 (USA, 2018). The temporal gradient holds within animals from the same genus, which would suggest that the translocated sequence is the oldest and the rest reflect more recent mutations. For the predicted secondary structures of the RNAs corresponding to the intragenomic rearrangement and adjacent sequences, the minimum free energy increases among variants from those with the most to those with the least similar sequence to the 5’-UTR insertion. Color scheme is as in previous figures.

There appears to be a temporal gradient with the most similar sequence (99%) in isolate KX964649 (China, 2011-10) to the least similar (59%) in isolate OK655840 (USA, 2018). The temporal gradient of decreasing similarity holds within rodents from the same genus, which would suggest that the translocated sequence is the oldest and the rest reflect more recent mutations. This is consistent with a possible common ancestor for all rodent α-CoVs sampled so far, with phylogenetic analyses suggesting relatively frequent host-jumping among the different rodent species (Tsoleridis et al. 2019). The minimum free energy of the predicted RNA secondary structures of the intragenomic rearrangement and adjacent sequences increases from the most similar to the least similar to the 5’-UTR insertion (Figure 9B). The function of the region of intragenomic rearrangement in nsp2 remains to be determined and it does not overlap with that contributing to inflammation via NF-kB activation in the α-CoV porcine transmissible gastroenteritis virus (Wang et al. 2018).

### Intragenomic rearrangements in N of bat **α**-CoVs subgenus Nyctacovirus

As shown in Figure 10A, in this intragenomic rearrangement in bat α-CoVs subgenus Nyctacovirus, a 115-nucleotide-long segment of the 5’-UTR is duplicated, inverted (negative-sense strand) and translocated to the proximal end of the nucleocapsid gene thereby encoding the first 38 amino acids of the amino-terminus of N. Other variants share the sequence with lesser similarity to the 5’-UTR sequence. There is a TRS-B sequence (AACUAA) at the beginning of the insertion, and the negative strand 5’-UTR-derived sequence also has a AACUAA sequence, which may have mediated the intragenomic rearrangement. The functional significance of this intragenomic rearrangement remains to be determined. Although in infectious bronchitis virus, the amino terminal domain of N has been shown to interact with nucleotide sequences in the 3’-UTR which is relevant to viral RNA packaging, the amino acids that are critical for such interaction are more distally located in the amino terminus (amino acids 76 or 94) (Tan et al. 2006; Zhou and Collisson, 2000) than those encoded by the intragenomic rearrangement in this case.

**Figure 10.**
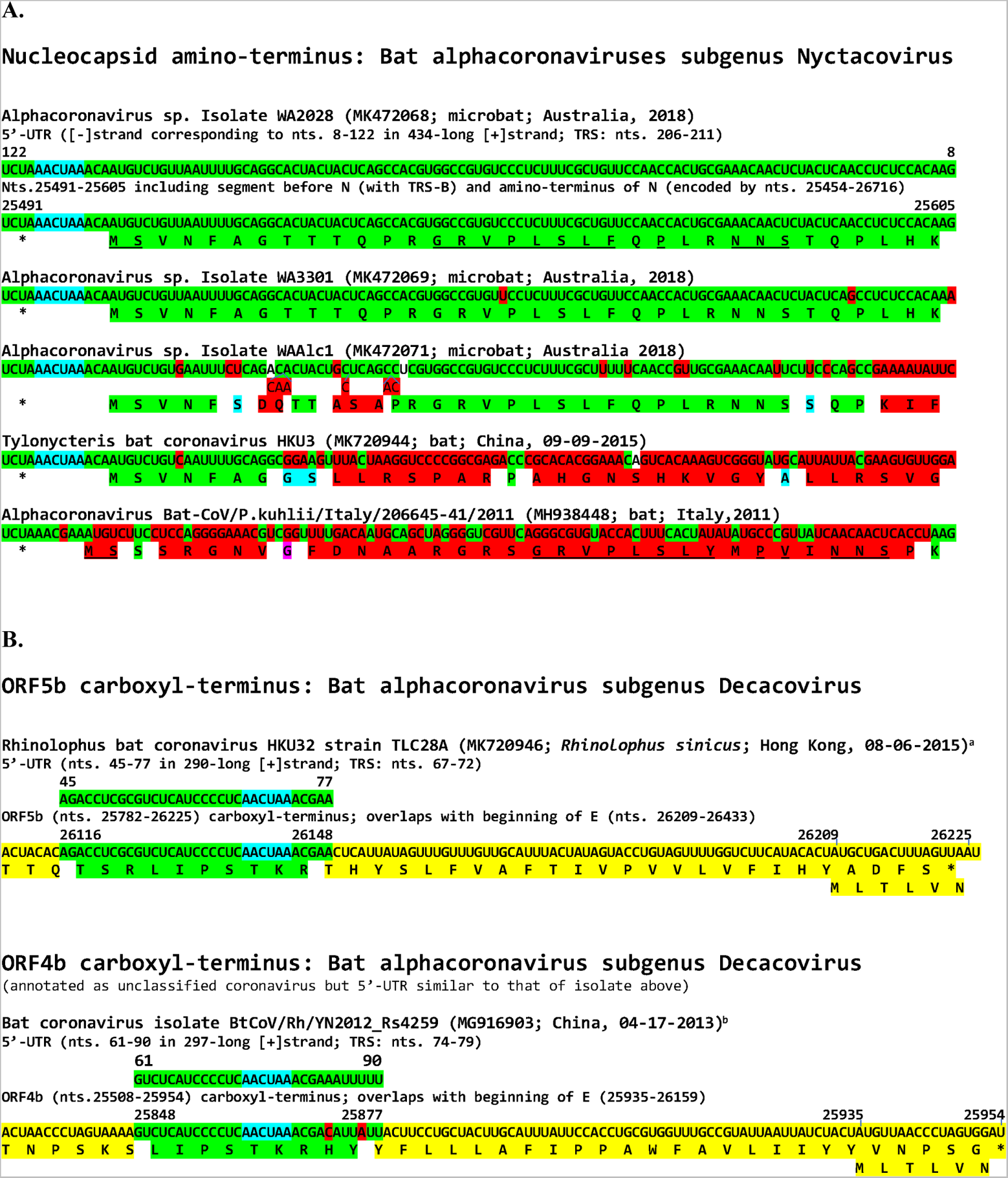
Intragenomic rearrangements in N of bat alphacoronaviruses subgenus Nyctacovirus. **(A) and in ORF5b or ORF4b of bat alphacoronaviruses subgenus Decacovirus.** Color scheme is as in Figure 9.

### Intragenomic rearrangements in ORF5b/4b of bat α-CoVs subgenus Decacovirus

Orb5b and ORF4b proteins are 53% similar (including conservative substitutions) between bat alphacoronaviruses subgenus Decacovirus shown in Figure 10B. In both sets of viruses, ORF5b or ORF4b overlap the beginning of the membrane (M) gene, i.e., there is no intergenic region between them and M. However, there is a TRS-B sequence (AACUAA) within the 3’ end of ORF5b and ORf4b where the intragenomic rearrangement occurs. Viruses with similar intragenomic rearrangements are for ORF5b: *Rhinolophus* bat coronavirus HKU32 strain TLC28A (MK720946), *Rhinolophus* bat coronavirus HKU32 strain TLC26A (MK720945; Hong Kong, 08-06-2005), Alphacoronavirus sp. strain bat/Yunnan/HcYN26/2020 (MZ081384; *Hipposideros cineraceus*; China, 07-29-2020), Alphacoronavirus sp. strain bat/Yunnan/RsYN12/2019 (MZ081386; *Rhinolophus sinicus*; China, 10-22-2019), Alphacoronavirus sp. strain bat/Yunnan/MmYN16/2020 (MZ081385; *Myotis muricola*; China, 04-18-2020), Alphacoronavirus sp. strain bat/Yunnan/RmYN21/2020 (MZ081387; *Rinolophus malayanus*; China, 06-03-2020); and for ORF4b: Bat coronavirus isolate BtCoV/Rh/YN2012_Rs4259 (MG916903; China, 04-17-2013), Bat coronavirus isolate BtCoV/Rh/YN2012_Rs4125 (MG916902; China, 09-16-2012). The functional significance of this intragenomic rearrangement remains to be determined.

### Intragenomic rearrangements derived from 5’-UTR sequences were not detected in some β-or α-, or in any γ- and δ-CoVs, and no 3’-UTR-derived intragenomic rearrangements were detected in any coronavirus

A listing of the other coronaviruses analyzed beyond the ones found to have intragenomic rearrangements is provided at the end of the Supplemental section. The directionality of translocation appears to be in the 5’ to 3’ direction as further underscored by the absence of 3’-UTR-derived insertions in any of the viruses analyzed.

## Discussion

We here describe intragenomic rearrangements involving 5’-UTR-derived sequences and the coding section of the genome of β- and α-CoVs. Figure 11A summarizes the locations of insertions (yellow arrows) in accessory, structural, and nonstructural genes of SARS-CoV-2, which for at least the accessory and structural genes appear to involve and/or affect the template switching mechanism by creating new regions of homology for interaction with TRS-L. We had previously reported (Patarca and Haseltine 2022) on the presence of conserved complementary sequences (CCSs) in the 5’- and 3’-UTRs potentially involved in circularization of the genome during subgenomic RNA synthesis. As shown in Figure 11B, the 5’-UTR-derived sequences involved in intragenomic rearrangements in SARS-CoV-2 shown in the present work usually include the TRS-L and span approximately half of the 5’ CCS, thus potentially facilitating circularization of the genome from locations closer to the 3’-UTR.

**Figure 11.**
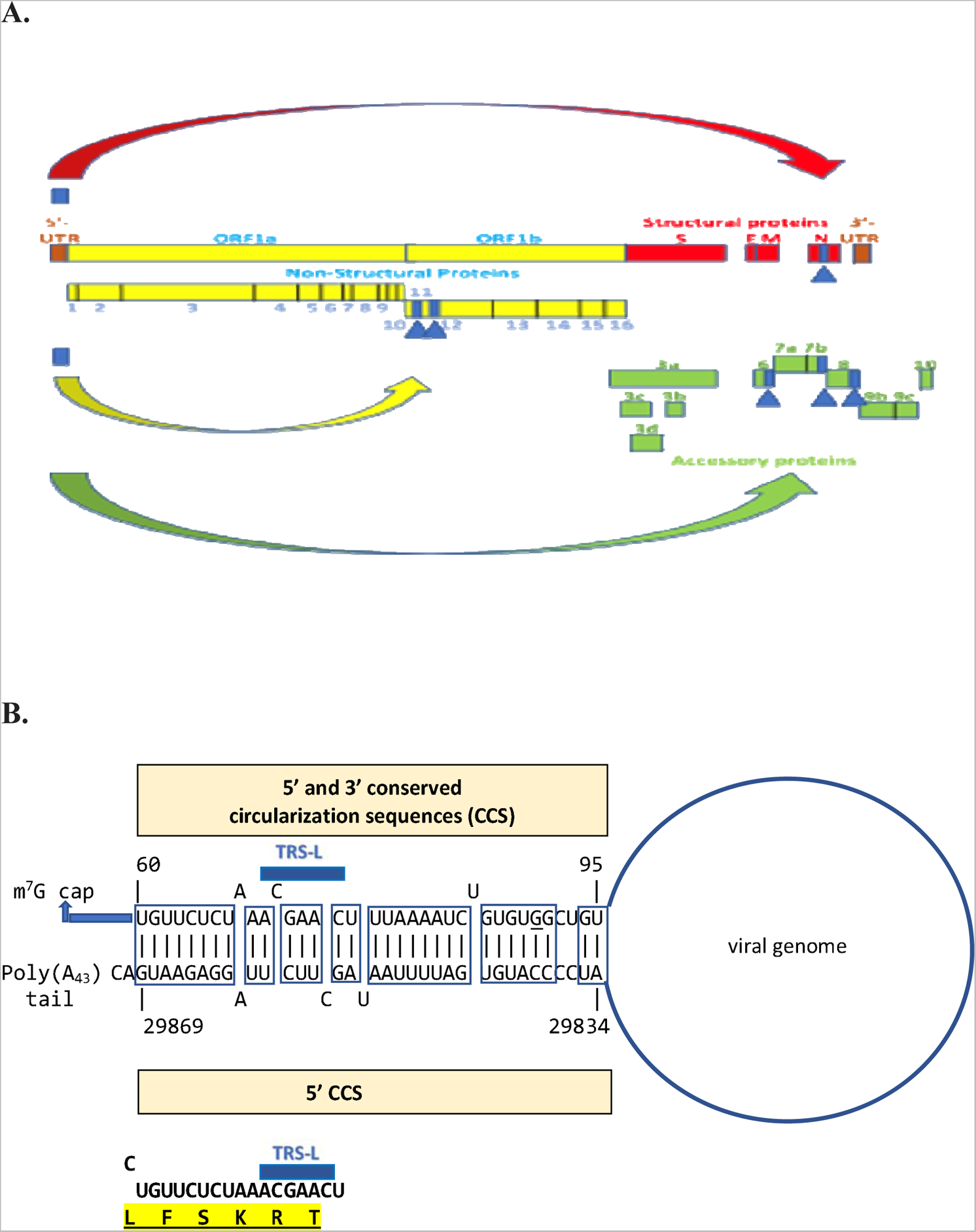
**A. Summary of locations of intragenomic rearrangements identified in SARS-CoV-2.** Nonstructural structural and accessory genes are represented in yellow, red, and green, respectively. **B. Partial overlap between the 5’-UTR nucleotide segment that is translocated to viral genes and a predicted sequence potentially involved in circularization of genome during viral replication.** The previously reported conserved complementary sequences (CCSs) in the 5’- and 3’ UTRs potentially involved in circularization of the genome during subgenomic RNA synthesis (Patarca and Haseltine 2022) are shown. The insertion sequences usually include the TRS-L and span approximately half of the 5’ CCS, thus potentially facilitating circularization of the genome from locations closer to the 3’-UTR.

Most of the 5’-UTR sequences duplicated and translocated include TRS-L. Extending the homology region of interaction between the TRS-L in the 5’-leader and the TRS-L introduced in a particular area of the body of the genome optimizes free minimum energy of the interaction. Such facilitation may favor expression of certain genes over that of others, thereby altering the hierarchy in gene expression. Because insertions are in various locations of viral genes, including some encoding nonstructural proteins, they may propitiate formation of new subgenomic RNAs thereby expanding the repertoire of proteins and even transforming noncanonical subgenomic messenger RNAs, i.e., not associated with TRS homology, to canonical ones. SARS-CoV-2 and other CoVs (Bentley et al. 2013) have been reported to generate noncanonical subgenomic RNAs in abundance, accounting for up to a third of subgenomic messenger RNAs in cell culture models of infection and increasing in proportion over time (Nomburg et al. 2020).

The structural genes control genome dissemination (Lauber et al. 2013) while the accessory genes in the same region of the genome may be involved in adaptation to specific hosts, modulation of the interferon signaling pathways, the production of pro-inflammatory cytokines, or the induction of apoptosis (Cui et al. 2021; Fang et al. 2021). Gaining insight into the effect of the amino acid changes introduced by the 5’-UTR-derived sequences is likely to shed light into pathogenesis and immune evasion mechanisms. For instance, a few point mutations can have a profound effect as exemplified by the few mutations in the C-terminus of the spike protein that transform the feline CoV associated with mild disease to one, the feline infectious peritonitis virus, that is generally lethal (Rottier et al. 2005).

Intragenomic rearrangements are yet another example of the tremendous genomic flexibility of coronaviruses which underlies changes in transmissibility, immune escape and/or virulence documented during the SARS-CoV-2 pandemic.

## Limitations

The intragenomic rearrangements involving 5’-UTR sequences were detected in all subgenera of β-coronaviruses infecting humans (i.e., Sarbecovirus, Embecovirus, and Merbecovirus) and in the Nobecovirus but not the Hibecovirus subgenera of CoVs infecting bats. There were only 3 Hibecovirus genomes in the database, which may account for the lack of detection of internal rearrangements in this subgenus most closely related to Sarbecoviruses. In this respect, the most diverse detection of

rearrangements in SARS-CoV-2 may reflect the bias generated by the presence in GenBank of SARS-CoV-2 isolates in up to 5 orders of magnitude greater number than any other CoV. However, the relative paucity of α-, γ-, or δ-CoV sequences available also applies to those of β-CoVs other than SARS-CoV-2 for which 5’-UTR rearrangements were found in notable proportions. Moreover, the present analysis included CoVs involved in large outbreaks such as the swine enteric CoVs of the α and δ genera and avian infectious bronchitis virus of the γ genus that have been studied over decades with hundreds of isolates characterized without apparent evidence for intragenomic rearrangements. The apparent absence of internal rearrangements in the latter viruses bodes well for the specificity of the findings described here for 4 of 5 subgenera of β-CoVs and 3 of 12 subgenera of α-CoVs.

Many sequences in the databases have incomplete 5’-UTRs rendering it difficult to comprehensively analyze them and to calculate more reliable proportions of variations. There are also partial genome and protein sequences, and we excluded sequences with undetermined amino acids. Nonetheless, for SARS-CoV-2, the frequency of variants with full-length insertions appears low relative to those with subsegments or other mutations relative to the reference strain in the same insertion area. One could posit that for hCoV-OC43 and hCoV-HKU-1, the apparently much higher frequency of intragenomic rearrangements involving 5’-UTR sequences might be driven by characterization of a greater number of isolates during epidemics with rearrangements possibly providing transmissibility, immune evasion and/or virulence advantages.

A limitation of the methods used for detecting these isolates is that they may not be viable, i.e., they may be associated with molecular diagnostic detection of virus but not necessarily culture conversion, or may represent artifacts of sequencing; however, their prevalence with redundancy in various locations and processing laboratories would be consistent with human-to-human transmission. Moreover, Turakhia et al. (2020), among others, have pointed out that systematic errors associated with lab-or protocol-specific practices affect some sequences in the repositories, which are predominantly or exclusively from single labs, co-localize with commonly used primer binding sites and are more likely to affect the protein-coding sequences than other similarly recurrent mutations. Although we cannot rule out that such systematic errors may underlie some if not all the rearrangements detected, the possibility is rendered less likely by the geographic and temporal diversity of the isolates with each intragenomic rearrangement (as underscored by the data in the Supplemental section), their presence in diverse variants of concern, as well as the occurrence of rearrangements in sequences from before the pandemic era and among diverse viruses of 2 genera and various subgenera in at least three hosts (humans, bats, and rodents). Moreover, it is unlikely that the insertion in the nucleocapsid gene of SARS-CoV-2 which encodes for a common co-mutation of adjacent sites that has been shown experimentally to have functional significance reflects an artifactual event.

Intragenomic rearrangements might be more common in isolates with large deletions, as exemplified by those involving the ORF6 (Tse et al. 2021), ORF7b and ORF8 of SARS-CoV-2, which in all cases affect the C-termini of the encoded proteins. The length of the insertion does not notably affect that of the protein in isolates without major genomic deletions. For 5’-UTR segments within viral genes, such as the examples shown in N, nsp12 and nsp3, or intergenic regions, the length of the protein or intergenic region appears not to be affected.

Variation driven by internal rearrangements is distinct from the non-homologous recombination events proposed as origins of Sarbecovirus/Hibecovirus/Nobecovirus β-CoV ORF3a by gene duplication followed by rapid divergence from M (Nikolaidis et al. 2021; Ouzounis et al. 2020) or of SARS-CoV-2 ORF8 from ORF7a (Neches et al. 2021). The mechanisms underlying intragenomic rearrangements warrant further study. Understanding the variation that they introduce also is of relevance in the design of prophylactic and therapeutic interventions for all coronaviruses, including a pan-β-coronavirus vaccine.

## Materials and methods

To assess the presence of 5’-UTR-derived insertions in the body of the genome, we used 5- to 10-amino acid stretches from the 3 reading frames of the translated 5’-UTR nucleotide sequence of SARS-CoV-2 (Wuhan reference, NC_045512) as query sequences to search the GenBank^®^ database using the Basic Alignment Search Tool (BLAST)P^®^ (<u>Protein BLAST: search protein databases using a protein query</u> <u>(nih.gov)</u>; Altschul et al. 1997) for SARS-CoV-2 and SARS-CoV-related viral proteins encoding similar stretches. All nonredundant translated CDS + PDB + SwissProt + PRF excluding environmental samples from WGS projects were searched specifying severe acute respiratory syndrome coronavirus 2 as organism.

Using the accession number listed in PubMed (<u>SARS-CoV-2 Resources - NCBI (nih.gov</u>) for the viral protein sequence, we obtained the respective nucleotide sequence and translated it using the insilico (<u>DNA to protein translation (ehu.es)</u> [Bikandy et al. 2004] and Expasy (<u>ExPASy - Translate tool</u> [Duvaud et al. 2021]) tools to determine by manual inspection and the BLASTN program (Johnson et al. 2008) if the nucleotide sequences encoding said stretches were identical to those in the 5’-UTR nucleotide sequence of SARS-CoV-2 or SARS-CoV-related viruses.

To detect isolates with similar insertions whose sequences had not been included in GenBank, we then searched the Global Initiative on Sharing All Influenza <u>Data (</u>GISAID) EpiFlu™ database of SARS-CoV-2 sequences (<u>GISAID - Initiative</u>; Elbe et al. 2017; Khare et al. 2021; Shu et al. 2017) using as queries the nucleotide sequences of the insertions plus adjoining 20 nucleotides on either side from the viral genomes. This approach is limited by the fact that maximum number of search results in GISAID is 30. Information on location and timing of isolate collection was obtained from the GenBank and GISAID databases.

We used the Rfam database (http://rfam.xfam.org/covid-19) with the curated Stockholm files containing UTR sequences, alignments and consensus RNA secondary structures of major genera of Coronoviridae; the representative RefSeq sequences for each genus obtained from the International Committee on Taxonomy of Viruses (ICTV) taxonomy Coronaviridae Study Group (2020 release; https://talk.ictvonline.org/ictv-reports/ictv_9th_report/positive-sense-rna-viruses-2011/w/posrna_viruses/223/coronaviridae-figures); the reference sequences in the GenBank database; and listings in publications involving phylogenetic analyses of alpha-, delta-, and gamma-coronaviruses from NCBI Taxonomy [Nikolaidis M. et al. 2021; Tsoleridis T. et al. 2019] to derive the 5’-UTRs of various CoVs.

We then utilized the 5’-UTR segments as query sequences to search for insertions in their respective genomes (nucleotide collection [nr/nt]; expect threshold: 0.05; mismatch scores: 2, −3; gap costs: linear). The GSAID database does not include sequences of CoVs other than SARS-CoV-2 and therefore could not be used for this analysis. Using nucleotide sequences instead of translated amino acid sequences from the 5’-UTR in the 3 reading frames as query sequences was unproductive to detect insertions in SARS-CoV-2 because of the large number of SARS-CoV-2 sequences in the GenBank database and the limit of 5000 results in the BLAST algorithm settings which yielded solely 5’-UTR sequences.

If the intragenomic rearrangement detected using the 5’-UTR sequences involved a coding region, we translated the 5’-UTR insertion and adjacent segments using the insilico (<u>DNA to protein translation</u> <u>(ehu.es)</u> [Bikandy et al. 2004] and Expasy (<u>ExPASy - Translate tool</u> [Duvaud et al. 2021]) tools.

RNA secondary structures of the 5’-UTR-derived insertion and adjacent sequences of the intragenomic rearrangement were visualized using forna, a force directed graph layout (ViennaRNA Web services; [Kerpedjiev et al. 2015]), and the optimal secondary structures and their minimal free energies were determined using the RNAfold webserver [Gruber et al. 2008; Lorenz et al. 2011].

In terms of the locations of the insertions in the body of the genomes, the boundaries of nonstructural, structural, and accessory open reading frames were determined based on GenBank annotation and from manual inspection of multiple alignments and sequence similarities. In the results presented, we excluded matches to entries corresponding to the 5’-leader sequences in mRNAs from full viruses or defective interfering RNA particles, as well as protein sequences with >80% unknown amino acids (represented by the letter X) in GenBank. We also tested the 3’-UTR sequences using the same approaches described for the 5’-UTR ones. The Supplementary section includes the accession numbers and collection site and date, and in some cases the SARS-CoV-2 lineages, for the isolates with intragenomic rearrangements involving 5’-UTR sequences.

## References

Alhammad YMO, Kashipathy MM, Roy A, et al. The SARS-CoV-2 Conserved Macrodomain Is a Mono-ADP-Ribosylhydrolase. J Virol. 2021;95(3):e01969–20.

Altschul SF, Madden TL, Schäffer AA, et al. Gapped BLAST and PSI-BLAST: a new generation of protein database search programs. Nucleic Acids Res. 1997;25:3389–3402.

Amoutzias GD, Nikolaidis M, Tryfonopoulou E, Chlichlia K, Markoulatos P, Oliver SG. The remarkable evolutionary plasticity of coronaviruses by mutation and recombination: Insights for the COVID-19 pandemic and the future evolutionary paths of SARS-CoV-2. Viruses. 2022;14:78.

Andersen KG, Rambaut A, Lipkin WI, Holmes EC, Garry RF. The proximal origin of SARS-CoV-2. Nat. Med. 2020;26(4):450–452.

Angelini MM, Akhlaghpour M, Neuman BW, Buchmeier MJ. Severe acute respiratory syndrome coronavirus nonstructural proteins 3, 4, and 6 induce double-membrane vesicles. MBio. 2013;4(4): e00524–13.

Beidas M, Chehadeh W. Effect of Human Coronavirus OC43 Structural and Accessory Proteins on the Transcriptional Activation of Antiviral Response Elements. Intervirology. 2018;61(1):30–35.

Beidas M, Chehadeh W. PCR array profiling of antiviral genes in human embryonic kidney cells expressing human coronavirus OC43 structural and accessory proteins. Arch Virol. 2018;163:2065–2072.

Bentley K, Keep SM, Armesto M, Britton P. Identification of a Noncanonically Transcribed Subgenomic mRNA of Infectious Bronchitis Virus and Other Gammacoronaviruses. J Virol. 2013; 87:2128–2136.

Bikandi, J., San Millán, R., Rementeria, A., and Garaizar, J. *In silico* analysis of complete bacterial genomes: PCR, AFLP-PCR, and endonuclease restriction. Bioinformatics. 2004;20:798–799.

Bobay L-M, O’Donnell AC, Ochman H. Recombination events are concentrated in the sspike protein region of betacoronaviruses. PLoS Genet. 2020;16:e1009272.

Boni MF, Lemey P, Jiang X, et al. Evolutionary origins of the SARS-CoV-2 sarbecovirus lineage responsible for the COVID-19 pandemic. Nat Microbiol. 2020;5:1408–1417.

Carlson CR, Asfaha JB, Ghent CM, et al. Phosphoregulation of Phase Separation by the SARS-CoV-2 N Protein Suggests a Biophysical Basis for its Dual Functions. Mol Cell. 2020 Dec 17;80(6):1092–1103.e4.

Chen SC, Olsthoorn RCL. Group-specific structural features of the 5’-proximal sequences of coronavirus genomic RNAs. Virology. 2010;401(1):29–41.

Corman VM, Muth D, Niemeyer D, Drosten C. Hosts and sources of endemic human coronaviruses. Adv Virus Res. 2018;100:163–188.

Crook JM, Murphy I, Carter DP, et al. Metagenomic identification of a new sarbecovirus from horseshoe bats in Europe. Sci Rep. 2021;11:14723.

Cui J, Li F, Shi Z-L. Origin and evolution of pathogenic coronaviruses. Nat Rev Microbiol. 2019;17:181–192.

Decaro N, Mari V, Campolo M, et al. Recombinant canine coronaviruses related to transmissible gastroenteritis virus of Swine and circulating in dogs. J Virol. 2009;83(3): 1532–1537.

Dudas G, Rambaut A. MERS CoV recombination: Implications about the reservoir and potential for adaptation. Virus Evol. 2016;2: vsv023.

Dutta NK, Mazumdar K, Gordy JT. The nucleocapsid protein of SARS–CoV-2: a target for vaccine development. J Virol. 2020;94(13): e00647–20.

Duvaud S, Gabella C, Lisacek F, Stockinger H, Ioannidis V, Durinx C. Expasy, the Swiss Bioinformatics Resource Portal, as designed by its user. Nucleic Acids Research. 2021. doi: 10.1093/nar/gks225.

Dwivedy A, Mariadasse R, Ahmad M, et al. Characterization of the NiRAN domain from RNA-dependent RNA polymerase provides insights into a potential therapeutic target against SARS-CoV-2. PLoS Comput Biol. 2021;17(9):e1009384.

Elbe, S. and Buckland-Merrett, G. Data, disease and diplomacy: GISAID’s innovative contribution to global health. Global Challenges. 2017;1:33–46.

Fang P, Fang L, Zhang H, Xia S, Xiao S. Functions of coronavirus accessory proteins: Overview of the state of the art. Viruses. 2021;13:1139.

Flower TG, Buffalo CZ, Hooy RM, Allaire M, Ren X, Hurley JH. Structure of SARS-CoV-2 ORF8, a rapidly evolving immune evasion protein. Proc Natl Acad Sci USA. 2021;118(2):e2021785118.

Forni D, Cagliani R, Clerici M, Sironi M. Molecular evolution of human coronavirus genomes. Trends Microbiol. 2017;25:35–48.

Forni D, Cagliani R, Sironi M. Recombination and positive selection differentially shaped the diversity of betacoronavirus subgenera. Viruses. 2020;12:1313.

Franco-Muñoz C, Álvarez-Díaz DA, Laiton-Donato K, et al. Substitutions in Spike and Nucleocapsid proteins of SARS-CoV-2 circulating in South America. Infect Genet Evol. 2020;85:104557.

Goldstein SA, Brown J, Pedersen BS, Quinlan AR, Elde NC. Extensive recombination-driven coronavirus diversification expands the pool of potential pandemic pathogens. bioRxiv, 2021.02.03.429646.

Gorbalenya AE, Baker SC, Baric RS, et al. The species Severe Acute Respiratory Syndrome-related coronavirus: Classifying 2019-NCoV and naming it SARS-CoV-2. Nat Microbiol. 2020;5:536–544.

Gordon DE, Hiatt J, Bouhaddou M, et al. Comparative host-coronavirus protein interaction networks reveal pan-viral disease mechanisms. Science. 2020;370(6521):eabe9403.

Graham RL, Baric RS. Recombination, reservoirs, and the modular spike. Mechanisms of coronavirus cross-species transmission. J Virol. 2010;84:3134–3146.

Graham RL, Deing DJ, Deming ME, et al. Evaluation of a recombination-resistant coronavirus as a broadly applicable, rapidly implementable vaccine platform. Commun Biol. 2018; 1(1): 1–10.

Grifoni A, Weiskopf D, Ramirez SI, et al. Targets of T Cell Responses to SARS-CoV-2 Coronavirus in Humans with COVID-19 Disease and Unexposed Individuals. Cell. 2020;181(7):1489–1501.e15.

Gruber AR, Lorenz R, Bernhart SH, Neuböck R, Hofacker IL. The Vienna RNA Websuite. Nucleic Acids Research. 2008; 36 (Issue suppl_2, 1 July): W70–W74. doi: 10.1093/nar/gkn188

Gussow AB, Auslander N, Faure G, Wolf YI, et al. Genomic determinants of pathogenicity in SARS-CoV-2 and other human coronaviruses. Proc Nat Acad Sci USA. 2020; 117(26):15193.

Hachim A, Kavian N, Cohen CA, et al. ORF8 and ORF3b antibodies are accurate serological markers of early and late SARS-CoV-2 infection. Nature immunology. 2020 Oct;21(10):1293–301.

Hagemeijer MC, Monastyrska I, Griffith J, van der Sluijs P, Voortman J, et al. Membrane rearrangements mediated by coronavirus nonstructural proteins 3 and 4. Virology. 2014;458:125–135.

Harrison, G. P., Mayo, M. S., Hunter, E., & Lever, A. M. (1998). Pausing of reverse transcriptase on retroviral RNA templates is influenced by secondary structures both 5’ and 3’ of the catalytic site. Nucleic acids research. 1998; 26(14):3433–3442.

Hartenian F, Nandakumar D, Lari A, Ly M, Tucker JM, Glausinger BA. The molecular virology of coronaviruses. J Biol Chem. 2020;295:12910–12934.

Hassan SS, Choudhury PP, Dayhoff GW 2nd, Aljabali AAA, et al. The importance of accessory protein variants in the pathogenicity of SARS-CoV-2. Arch Biochem Biophys. 2022;717:109124.

He R, Leeson A, Ballantine M, et al. Characterization of protein-protein interactions between the nucleocapsid protein and membrane protein of the SARS coronavirus. Virus Res. 2004;105(2):121–125.

ICTV Coronaviridae Study Group. International Committee on Taxonomy of Viruses (ICTV). 2021. Available from: https://talk.ictvonline.org/ictv-reports/ictv_9th_report/positive-sense-rna-viruses-2011/w/posrna_viruses/223/coronaviridae-figures.

Imbert I, Snijder EJ, Dimitrova M, Guillemot JC, Lécine P, Canard B. The SARS-Coronavirus PLnc domain of nsp3 as a replication/transcription scaffolding protein. Virus research. 2008; 133(2):136–148.

Islam MR, Hoque MN, Rahman MS, Alam ASMRU, Akther M, et al. Genome-wide analysis of SARS-CoV-2 virus strains circulating worldwide implicates heterogeneity. Sci Rep. 2020;10(1):14004.

Jackson B, Boni MF, Bull MJ, et al. Generation and transmission of interlineage recombinants in the SARS-CoV-2 pandemic. Cell. 2021 Sep 30;184(20):5179–5188.

Jaroszewski L, Iyer M, et al. The interplay of SARS-CoV-2 evolution and constraints imposed by the structure and functionality of its proteins. PLoS computational biology. 2021; 17(7):e1009147.

Johnson M, Zaretskaya I, Raytselis Y, Merezhuk Y, McGinnis S, Madden TL. NCBI BLAST: a better web interface. Nucleic Acids Res. 2008; 36(Web Server issue): W5–9. doi: 10.1093/nar/gkn201.

Johnson BA, Zhou Y, Lokugamage KG, et al. Nucleocapsid mutations in SARS-CoV-2 augment replication and pathogenesis. bioRxiv [Preprint]. 2021 Oct 15:2021. doi: 10.1101/2021.10.14.464390.

Kemp BE, Graves DJ, Benjamini E, Krebs EG. Role of multiple basic residues in determining the substrate specificity of cyclic AMP-dependent protein kinase. J Biol Chem. 1977;252(14):4888–4894.

Kennelly PJ, Krebs EG. Consensus sequences as substrate specificity determinants for protein kinases and protein phosphatases. J Biol Chem. 1991;266(24):15555–15558.

Kerpedjiev P, Hammer S, Hofacker IL. Forna (force-directed RNA): Simple and effective online RNA secondary structure diagrams. Bioinformatics. 2015; 31:3377–3379.

Khavinson V, Terekhov A, Kormilets D, Maryanovich A. Homology between SARS CoV-2 and human proteins. Sci Rep. 2021;11:17199. doi: 10.1038/s41598-021-96233-7.

Khare, S., et al. GISAID’s Role in Pandemic Response. China CDC Weekly. 2021;3(49): 1049–1051.

Latinne A, Hu B, Olival KJ, Zhu G, Zhang L, Li H, Chmura AA, Field HE, et al. Origin and cross-species transmission of bat coronaviruses in China. Nature Communications. 2020;11(1):1–5.

Lau SKP, Wong EYM, Tsang CC, et al. Discovery and sequence analysis of four deltacoronaviruses from birds in the Middle East reveal interspecies jumping with recombination as a potential mechanism for avian-to-avian and avian-to-mammalian transmission. J Virol. 2018;92: e00265–18.

Lauber C, Goeman JJ, Parquet MDC, Thi Nga P, Snijder EJ, Morita K, Gorbalenya AE. The footprint of genome architecture in the largest genome expansion in RNA viruses. PLoS Pathogen. 2013;9:e1003500.

Lehmann KC, Gulyaeva A, Zevenhoven-Dobbe JC, et al. Discovery of an essential nucleotidylating activity associated with a newly delineated conserved domain in the RNA polymerase-containing protein of all nidoviruses. Nucleic Acids Res. 2015;43(17):8416–8434.

Lei J, Kusov Y, Hilgenfeld R. Nsp3 of coronaviruses: Structures and functions of a large multi-domain protein. Antiviral research. 2018;149:58–74.

Li JY, Liao CH, Wang Q, Tan YJ, Luo R, Qiu Y, Ge XY. The ORF6, ORF8 and nucleocapsid proteins of SARS-CoV-2 inhibit type I interferon signaling pathway. Virus research. 2020;286:198074.

Lin X, Fu B, Yin S, Li Z, Liu H, Zhang H, Xing N, Wang Y, Xue W, et al. ORF8 contributes to cytokine storm during SARS-CoV-2 infection by activating IL-17 pathway. iScience. 2021;24(4):102293.

Liu DX, Fung TS, Chong KK, Shukla A, Hilgenfeld R. Accessory proteins of SARS-CoV and other coronaviruses. Antiviral Res, 2014;109:97–109. doi: 10.1016/j.antiviral.2014.06.013.

Lo C-Y, Tsai T-L, Lin C-N, Lin C-H, Wu H-Y. 2019. Interaction of coronavirus nucleocapsid protein with the 5’- and 3’-ends of the coronavirus genome is involved in genome circularization and negative strand RNA synthesis. FEBS J. 2019;286:3222–3239.

Lorenz R, Bernhart SH, Höner zu Siederdissen C, Tafer H, Flamm C, Stadler PF, Hofacker IL. “ViennaRNA Package 2.0”, Algorithms for Molecular Biology. 2011; 6(1):26.

Lu S, Ye Q, Singh D, Cao Y, et al. The SARS-CoV-2 nucleocapsid phosphoprotein forms mutually exclusive condensates with RNA and the membrane-associated M protein. Nat Commun. 2021;12(1):502.

Lytras S, Hughes J, Martin D, de Klerk A, Lourens R, Kosakovsky Pond SL, Xia W, et al. Exploring the natural origins of SARS-CoV-2 in the light of recombination. Genome Biol Evol. 2022;evac018.

Madhugiri R, Karl N, Petersen D, Lamkiewicz K, et al. Structural and functional conservation of cis-acting RNA elements in coronavirus 5’-terminal genome regions. Virology. 2018;517:44–55.

Makino S, Keck JG, Stohlman SA, Lai MM. High-frequency RNA recombination of murine coronaviruses. J Virol. 1986;57:729–737.

Matthews KL, Coleman CM, van der Meer Y, Snijder EJ, Frieman MB. The ORF4b-encoded accessory proteins of Middle East respiratory syndrome coronavirus and two related bat coronaviruses localize to the nucleus and inhibit innate immune signalling. J Gen Virol. 2014;95(Pt 4):874.

McBride R, van Zyl M, Fielding BC. The coronavirus nucleocapsid is a multifunctional protein. Viruses. 2014;6(8):2991–3018. doi: 10.3390/v6082991.

Menachery, V. D., Yount, B. L., Jr, Debbink, K., et al. (2015). A SARS-like cluster of circulating bat coronaviruses shows potential for human emergence. Nature medicine. 2015;21(12):1508–1513.

Miao Z, Tidu A, Eriani G, Martin F. Secondary structure of the SARS-CoV-2 5’-UTR. RNA Biology. 2021;18(4):447–456.

Mounir S, Talbot PJ. Molecular characterization of the S protein gene of human coronavirus OC43. J Gen Virol. 1993;74:1981–1987.

Mourier, T., Shuaib, M., Hala, S. et al. SARS-CoV-2 genomes from Saudi Arabia implicate nucleocapsid mutations in host response and increased viral load. Nat Commun. 2022;13, 601. https://doi.org/10.1038/s41467-022-28287-8

Neches RY, Kyrpides NC, Ouzounis CA. Atypical divergence of SARS-CoV-2 Orf8 from Orf7a within the coronavirus lineage suggests potential stealthy viral strategies in immune evasion. Mbio. 2021;12(1):e03014–20.

Niemeyer D, Zillinger T, Muth D, et al. Middle East respiratory syndrome coronavirus accessory protein 4a is a type I interferon antagonist. J. Virol. 2013;87(22):12489–12495.

Nikolaidis M, Markoulatos P, van de Peer Y, Oliver SG, Amoutzias GD. The neighborhood of the spike gene is a hotspot for modular intertypic homologous and non-homologous recombination in coronavirus genomes. Mol Biol Evol. 2021;msab 292.

Nomburg J, Meyerson M, De Caprio JA. Pervasive generation of non-canonical subgenomic RNAs by SARS-CoV-2. Genome Medicine. 2020;12:108.

Oliveira SC, de Magalhães MTQ, Homan EJ. Immunoinformatic Analysis of SARS-CoV-2 Nucleocapsid Protein and Identification of COVID-19 Vaccine Targets. Front Immunol. 2020;11:587615.

Ouzounis C. A. A recent origin of Orf3a from M protein across the coronavirus lineage arising by sharp divergence. Computational and structural biotechnology journal. 2020;18:4093–4102.

Park GJ, Osinski A, Hernandez G, et al. The mechanism of RNA capping by SARS-CoV-2. bioRxiv 2022.02.07.479471; doi: https://doi.org/10.1101/2022.02.07.479471

Patarca R, Haseltine WA. Circularization via complementary sequences in the 5’ and 3’ termini may facilitate replication of SARS coronaviruses. Authorea. January 04, 2022. doi: 10.22541/au.164132044.46753705/v1

Pickering B, Lung O, Maguire F, et al. Highly divergent white-tailed deer SARS-CoV-2 with potential deer-to-human transmission. bioRxiv 2022.02.22.481551. doi: https://doi.org/10.1101/2022.02.22.481551

Pollett S, Conte MA, Sanborn M, Jarman RG, Lidl GM, et al. A comparative recombination of analysis of human coronaviruses and implications for the SARS-CoV-2 pandemic. Sci Rep. 2021;11:17365.

Pustovalova Y, Gorbatyuk O, Li Y, et al. Backbone and Ile, Leu, Val methyl group resonance assignment of CoV-Y domain of SARS-CoV-2 non-structural protein 3. Biomol NMR Assign. 2021;18:1–6.

Redondo N, Zaldívar-López S, Garrido JJ, and Montoya M. SARS-CoV-2 Accessory Proteins in Viral Pathogenesis: Knowns and Unknowns. Front. Immunol. 2021;12:708264

Rottier PJM, Nakamura K, Schellen P, Volders H, Hajema BJ. Acquisition of macrophage tropism during the pathogenesis of feline infectious peritonitis is determined by mutations in the feline coronavirus spike protein. J Virol. 2005;79:14122–14130.

Sawicki SG, Sawicki DL, Siddell SG. A contemporary view of coronavirus transcription. J Virol. 2007;81:20–29.

Shang J, Han N, Chen Z, Peng Y, Li L, Zhou H, Ji C, Meng J, Jiang T, Wu A. Compositional diversity and evolutionary pattern of coronavirus accessory proteins. Brief. Bioinform. 2021; 221267–1268.

Shu, Y. and McCauley, J. GISAID: from vision to reality. EuroSurveillance 2017;22(13) doi:10.2807/1560-7917.ES.2017.22.13.30494

Simon-Loriere E, Holmes EC. Why do RNA viruses recombine? Nat Rev Microbiol. 2011;9(8):617–626.

Siu KL, Yeung ML, Kok KH, Yuen KS, Kew C, et al. Middle east respiratory syndrome coronavirus 4a protein is a double-stranded RNA-binding protein that suppresses PACT-induced activation of RIG-I and MDA5 in the innate antiviral response. J Virol. 2014;88(9): 4866–4876.

Slanina H, Madhugiri R, Bylapudi G, Schultheiß K, Karl N, Gulyaeva A, Gorbalenya AE, Linne U, Ziebuhr J. Coronavirus replication-transcription complex: Vital and selective NMPylation of a conserved site in nsp9 by the NiRAN-RdRp subunit. Proc Natl Acad Sci USA. 2021;118(6):e2022310118.

Song H-D, Tu C-C, Zhang G-W, Wang S-Y, Su S, Wong G, Shi W, Liu J, Lai ACK, et al. Epidemiology, genetic recombination, and pathogenesis of coronaviruses. Trends Microbiol. 2016;24:490–502.

Stukalov A, Girault V, Grass V, Karayel O, Bergant V, Urban C, Haas DA, et al. Multilevel proteomics reveals host perturbations by SARS-CoV-2 and SARS-CoV. Nature. 2021;594(7862):246–252.

Surjit M, Kumar R, Mishra RN, Reddy MK, Chow VT, Lal SK. The severe acute respiratory syndrome coronavirus nucleocapsid protein is phosphorylated and localizes in the cytoplasm by 14-3-3-mediated translocation. J Virol. 2005; 79(17):11476–11486.

Tan YW, Fang S, Fan H, Lescar J, Liu DX. Amino acid residues critical for RNA-binding in the N-terminal domain of the nucleocapsid protein are essential determinants for the infectivity of coronavirus in cultured cells. Nucleic acids research, 2006; 34(17):4816–4825.

Temmam S, Vongphayloth K, Baquero Salazar E, et al. Coronaviruses with a SARS-CoV-2-like receptor-binding domain allowing ACE2-mediated entry into human cells isolated from bats of Indochinese peninsula. https://doi.org/10.21203/rs.3.rs-871965/v1 (2020).

Thorne LG, Bouhaddou M, Reuschl AK, Zuliani-Alvarez L, Polacco B, Pelin A, Batra J, Whelan M, et al. Evolution of enhanced innate immune evasion by SARS-CoV-2. Nature. 2022;602(7897), 487–495.

Tse H, Lung DC, Wong SC, et al. Emergence of a Severe Acute Respiratory Syndrome Coronavirus 2 Virus variant with novel genomic architecture in Hong Kong. Clin Infect Dis. 2021;73(9):1696–1699.

Tsoleridis T, Chappell JG, Onianwa O, Marston DA, Fooks AR, Monchatre-Leroy E, Umhang G, Müller MA, Drexler JF, Drosten C, Tarlinton, RE, McClure CP, Holmes EC, Ball JK. Shared Common Ancestry of Rodent Alphacoronaviruses Sampled Globally. Viruses. 2019; 11(2):125.

Tugaeva KV, Hawkins DEDP, Smith JLR, et al. The Mechanism of SARS-CoV-2 Nucleocapsid Protein Recognition by the Human 14-3-3 Proteins. J Mol Biol. 2021 Apr 16;433(8):166875.

Tung HYL, Limtung P. Mutations in the phosphorylation sites of SARS-CoV-2 encoded nucleocapsid protein and structure model of sequestration by protein 14-3-3. Biochem Biophys Res Comm. 2020;532:134–138.

Turakhia, Y., De Maio, N., Thornlow, B., Gozashti, L., Lanfear, R., Walker, C. R., Hinrichs, A. S., et al. Stability of SARS-CoV-2 phylogenies. PLoS genetics. 2020;16(11); e1009175.

Turakhia Y, Thornlow B, Hinrichs AS, McBroome J, Ayala N, et al. Pandemic-Scale phylogenomics reveals elevated recombination rates in the SARS-CoV-2 spike region. bioRxiv. 2021 Jan 1.

Ujike M, Taguchi F. Recent Progress in Torovirus Molecular Biology. Viruses. 2021, 13(3):435.

Valcarcel A, Bensussen A, Álvarez-Buylla ER, Díaz J. Structural Analysis of SARS-CoV-2 ORF8 Protein: Pathogenic and Therapeutic Implications. Front Genet. 2021; 12:693227.

VanInsberghe D, Neish AS, Lowen AC, Koelle K. Recombinant SARS-CoV-2 genomes are currently circulating at low levels. bioRxiv. 2021 Jan 1:2020–08.

Van Marle G, Luytjes W, Van der Most RG, et al. Regulation of coronavirus mRNA transcription. J Virol. 1995; 69(12):7851–7856.

Vijgen L, Keyaerts E, Moës E, Thoelen I, Wollants E, Lemey P, Vandamme A-M, Van Ranst M. Complete genomic sequence of human coronavirus OC43: Molecular clock analysis suggests a relatively recent zoonotic coronavirus transmission event. J Virol. 2005;79:1595–1604.

Wang L., Qiao, X., Zhang, S., Qin, Y., Guo, T., Hao, Z., Sun, L., Wang, X., Wang, Y., Jiang, Y., Tang, L., Xu, Y., & Li, Y. (2018). Porcine transmissible gastroenteritis virus nonstructural protein 2 contributes to inflammation via NF-κ B activation. Virulence, 9(1), 1685–1698.

Wang X, Lam JY, Wong WM, Yuen CK, Cai JP, Au SW, Chan JF, et al. Accurate Diagnosis of COVID-19 by a Novel Immunogenic Secreted SARS-CoV-2 orf8 Protein. mBio. 2020 Oct 20;11(5):e02431–20.

Wang D, Jiang A, Feng J, et al. The SARS-CoV-2 subgenome landscape and its novel regulatory features. Molecular Cell. 2021;81:2135–2147.

Weiss SR, Navas-Martin S. Coronavirus pathogenesis and the emerging pathogen severe acute respiratory syndrome coronavirus. Microbiology and molecular biology reviews. 2005;69(4):635–664.

Wille M, Holmes EC. Wild birds as reservoirs for diverse and abundant gamma- and deltacoronaviruses. FEMS microbiology reviews. 2020; 44(5):631–644.

Wong AC, Li X, Lau SK, Woo PC. Global epidemiology of bat coronaviruses. Viruses. 2019;11(2):174.

Woo PC, Lau SK, Huang Y, Yuen KY. Coronavirus diversity, phylogeny and interspecies jumping. Experimental Biology and medicine. 2009;234(10):1117–27.

Wu H, Xing N, Meng K, Fu B, Xue W, Dong P, et al. Nucleocapsid mutations R203K/G204R increase the infectivity, fitness, and virulence of SARS-CoV-2. Cell Host & Microbe. 2021;29:1788–1801.

Yan L, Ge J, Zheng L, Zhang Y, et al. Cryo-EM Structure of an Extended SARS-CoV-2 Replication and Transcription Complex Reveals an Intermediate State in Cap Synthesis. Cell. 2021;184(1):184–193.e10.

Yang R, Zhao Q, Rao J, Zeng F, Yuan S, Ji M, Sun X, et al. SARS-CoV-2 Accessory Protein ORF7b Mediates Tumor Necrosis Factor-α-Induced Apoptosis in Cells. Front Microbiol. 2021;12:654709.

Yang Y, Zhang L, Geng H, et al. The structural and accessory proteins M, ORF 4a, ORF 4b, and ORF 5 of Middle East respiratory syndrome coronavirus (MERS-CoV) are potent interferon antagonists. Protein & cell. 2013;4(12):951–961.

Yang Y, Yan W, Hall AB, Jiang X. Characterizing transcriptional regulatory sequences in coronaviruses and their role in recombination. Mol Biol Evol. 2021;38:1241–1248.

Yao H, Song Y, Chen Y, et al. Molecular Architecture of the SARS-CoV-2 Virus. Cell. 2020;183(3):730–738.e13. doi: 10.1016/j.cell.2020.09.018.

Zavadil J, Bitzer M, Liang D, et al. Genetic programs of epithelial cell plasticity directed by transforming growth factor-β. Proceedings of the National Academy of Sciences. 2001;98(12):6686–6691.

Zhang X, Liao C-L, Lai M. Coronavirus leader RNA regulates and initiates subgenomic mRNA transcription both in trans and in cis. J Virol. 1994; 8(8):4738–4746.

Zhang Y, Chen Y, Li Y, et al. The ORF8 protein of SARS-CoV-2 mediates immune evasion through down-regulating MHC-Ι. Proc Natl Acad Sci USA. 2021;118(23):e2024202118.

Zhou M, Collisson EW. The amino and carboxyl domains of the infectious bronchitis virus nucleocapsid protein interact with 3’ genomic RNA. Virus research. 2000; 67(1):31–39.

