## Supplemental data for "Intragenomic rearrangements in SARS-CoV-2, other betacoronaviruses, and alphacoronaviruses"

**Supplemental legend to Figure 2 in main text**

**SARS-CoV-2 variants (collection site and date in parentheses) with 5’-UTR-derived insertions modifying the ORF8 carboxyl terminus; those shown in Figure 2 in main text are highlighted**

**a.** NC_045512 (Wuhan reference); QUP34336 (USA/Minnesota, 2021-04-05); **b.** QVJ62740 (USA/Michigan, 2021-04-13), QUQ07869 (USA/Florida, 2021-04-04), QTM63997 (USA/Alabama, 2020-10-03), QTZ79380 (USA/Michigan, 2021-03-16), UCP53601 (USA/Arizona, 2020-10-29), QUA38826 (USA/Ohio, 2021-03-09), QTY87294 (USA/New Jersey, 2021-03-18), QTX61599 (USA/Michigan, 2021-02-24), QUA76009 (USA/Illinois, 2021-04-05), QUB00435 (USA/California, 2021-04-011), QVX18739 (USA/Minnesota, 2021-04-16), QVX54931 (USA/Virginia, 2021-05-07), QTW56832 (USA/Michigan, 2021-03-28), QTX80560 (USA/Michigan, 2021-03-08), QSV23506 (USA/Massachusetts, 2021-02-20), QTW56196 (USA/Michigan, 2021-03-28), UBZ78873 (USA/Virginia, 2021-08-15), QTM32955 (USA/California, 2021-03-13), QUP09826 (USA/Michigan, 2021-04-01), QVO97167 (USA/Michigan, 2021-03-10), QTY99266 (USA/Michigan, 2021-03-18), QQE72547 (USA/Florida, 2021-11-17), QUD47078 (USA/Michigan, 2021-04-03), UAZ71320 (USA/Michigan, 2021-03-22), UAP73316 (USA/Texas, 2021-08-20), UER42693 (USA/California, 2021-09-25), UBN91115 (USA/Florida, 2021-03-19), UCZ40652 (USA/Texas, 2021-10-03), QVM43844 (USA/Michigan, 2021-05-02), QZQ76855 (USA/Florida, 2021-05-24), QTB05507 (USA/Arkansas, 2021-03-02), QXG11190 (USA/Kentucky, 2021-05-24), QWA62675 (USA/Florida, 2021-05-08), QTZ60073 (USA/Pennsylvania, 2021-01-23); QUD47078 (USA/Michigan, 2021-04-03); **c.** QTZ60073 (USA/Pennsylvania, 2021-01-23),UBR92989 (USA/California, 2021-07-28), QTW56256 (USA/Pennsylvania, 2021-03-28), QTB05507 (USA/Arkansas, 2021-03-02), QTY29701 (USA/New York, 2021-04-02), QTP28777 (USA/South Carolina, 2021-03-15), QTC77450 (USA/Florida, 2021-02-21), UAU97712 (USA/Kansas, 2021-08-10), QUQ11804 (UA/Georgia, 2021-03-31), UFB90828 (USA/Massachusetts, 2021-10-27), QUC77139 (USA/New York, 2021-03-26), QSG76485 (USA/Texas, 2021-02-13), QRY28882 (USA/Washington, 2021-11-14), QUV66274 (USA/Washington, 2021-02-25), QUE36350 (USA/New Jersey, 2021-04-02), QZF56095 (USA/Nevada, 2021-07-18), QTG51220 (USA/Alabama, 2021-03-09), QTC29731 (USA/Washington, 2021-02-26), QUG22146 (USA/Arizona, 2021-02-03), QTM53955 (USA/California, 2021-03-11), UBG95816 (USA/Texas, 2021-05-27), QUA27428 (USA/New York, 2021-02-26), QYN93853 (USA/Minnesota, 2021-03-16), QWO65210 (USA/Illinois, 2021-05-26), QSW55172 (USA/California, 2021-12-18), QZC48693 (USA/California, 2021-01-08), QTZ59919 (USA/North Carolina, 2021-01-24), UEZ74290 (USA/Florida, 2021-06-13), QVU83077 (USA/West Virginia, 2021-05-03), UCP75589 (USA/Arizona, 2021-09-16), QSL79713 (USA/West Virginia, 2021-02-10), QQV24415 (USA/Washington, 2021-11-03), UDB06943 (USA/California, 2021-01-21), QWU51722 (USA/Missouri, 2021-03-31), QTP78406 (USA/Arizona, 2021-03-22; QWA62675 (USA/Florida, 2021-05-08); **d.** UAP73316 (USA/Texas, 2021-08-20), UER42693 (USA/California, 2020-09-25); UCZ40652 (USA/Texas, 2021-10-03); **e.** QXY14087 (USA/California, 2021-01-13), QUC83315 (USA/Michigan, 2021-03-26), QQZ32498 (USA/California, 2021-01-08), QQA02853 (USA/Maryland, 2020-11-11), QPG83352 (USA/Florida, 2020-05-29), QRV12235 (USA/Maryland, 2021-01-25), UEK15624 (USA/Colorado, 2020-09-17), QQA02817 (USA/Maryland, 2020-11-11), UCP62018 (USA/Arizona, 2020-10-03), UEK18017 (USA/Colorado, 2020-09-22), UEH39369 (USA/California, 2021-09-28), QZJ67737 (USA/Arizona, 2020-08-03), UFK15561 (USA/Minnesota, 2021-11-05), UFB92196 (USA/North Carolina, 2021-10-27), QZP72779 (USA/Alabama, 2021-07-30), QSX87276 (USA/Pennsylvania, 2021-02-25), UEU99280 (USA/Pennsylvania, 2021-11-06), UEL99603 (USA/California, 2021-10-12), UEU89370 (USA/Minnesota, 2021-11-02), QRG28603 (USA/Florida, 2020-09-05), QUF21578 (USA/Texas, 2021-03-27), UEN39474 (USA/Illinois, 2021-10-10), QUA15764 (USA/Texas, 2021-01-10), QUA44810 (USA/Pennsylvania, 2021-03-15), QVU27766 (USA/North Carolina, 2021-05-01), QUA44621 (USA/North Carolina, 2021-03-14), QYX84583 (USA/New Hampshire, 2021-03-10), QZA59779 (USA/Pennsylvania, 2020-12-03), QSG76473 (USA/Massachusetts, 2021-02-11), QUA13851 (USA/Missouri, 2020-12-06), QPG83362 (USA/Florida, 2020-05-29), QYV28981 (USA/New Jersey, 2021-02-26), QSF03724 (USA/Texas, 2021-02-07), QSL82762 (USA/Pennsylvania, 2021-02-13), QSX25919 (USA/South Carolina, 2021-02-03), QNU10627(USA/Florida, 2021-06-02), QTW58176 (USA/Virginia, 2021-03-29), QZC48492 (USA, California, 2021-01-11), QYM26447 (USA/California, 2020-12-23), QUR38225 (USA/New Jersey, 2021-04-12), QTZ61745 (USA/Arkansas, 2021-02-05), UBX59703 (USA/Minnesota, 2021-09-14), UDA69418 (USA/Virginia, 2021-04-07), QSX71636 (USA/South Carolina, 2021-01-22), QTC08790 (USA/South Carolina, 2021-02-24), QUA17514 (USA/West Virginia, 2021-01-25), UEK12552 (USA/Colorado, 2020-09-22), QSM30820 (USA/California, 2020-12-12-16), QTK20565 (USA/Michigan, 2021-03-16), QSN85839 (USA/Florida, 2021-02-10), QQX23234 (USA/Texas, 2020-10-02), QSF07756 (USA/Pennsylvania, 2021-01-29), QTC79814 (USA/Pennsylvania, 2021-02-22), QQV30431 (USA/California, 2021-01-07), UBE04273 (USA/Tennessee, 2021-09-02), QNL24003 (USA/Maryland, 2020-07-12), QUA19121 (USA/Illinois, 2021-01-29), QSF03424 (USA/California, 2021-02-07), QSO01007 (USA/New Jersey, 2021-02-10), QWO77958 (USA/Utah, 2021-01-26), QZM76691 (USA/New Mexico, 2021-07-19), UCR39596 (USA/Florida, 2021-09-10), UAA61445 (USA/Minnesota, 2021-08-14), QUC86568 (USA/New Hampshire, 2021-03-24), QND77013 (USA/Maine, 2020-03-20), QQN91746 (USA/California, 2020-12-26), UFL93464 (USA/Colorado, 2020-06-15), QXY46820 (USA, New York, 2021-03-23), QZC52177 (USA/California, 2020-07-17), QUH70396 (USA/Michigan, 2021-04-05), QYM30783 (USA/California, 2020-12-08), UET28764 (USA/Pennsylvania, 2021-10-23), QKE12265 (USA/Florida, 2020-04-20), QSX90861 (USA/Massachusetts, 2021-01-12), QUE08549 (UA/Minnesota, 2021-03-28), QSO13388 (USA/Kansas, 2021-02-14), QQH16681 (Pakistan, 2020-04-16), QSJ40506 (USA/Georgia, 2021-02-11), QXI74936 (USA/Arizona, 2021-01-04), UDE63363 (USA/West Virginia, 2021-09-20), QTZ59862 (USA/Florida, 2021-01-24), QQP31916 (USA/Maryland, 2020-12-30), UAJ33375 (USA/California, 2021-08-04), QSX88451 (USA/Pennsylvania, 2021-02-26), QTA53245 (USA/Indiana, 2021-02-04), QSS80876 (USA/Texas, 2021-02-22), QUA15569 (USA/Florida, 2021-01-04), QQN92800 (USA/Tennessee, 2020-11-25), UCO66724 (USA/Indiana, 2021-09-15), QRX62333 (USA/Maryland, 2021-01-31), QTS06851 (USA/Wisconsin, 2021-03-21), QTD08121 (USA/Virginia, 2021-02-28), QWE67901 (USA/District of Columbia, 2021-05-15), QQL13208 (USA/Maryland, 2020-12-01), QVU02466 (USA/Michigan, 2021-04-15), UCN24542 (Kenya, 2020-11-17), UCR20446 (USA/Georgia, 2021-09-03), UCC02971 (USA/New York, 2021-08-26), QTP30077 (USA/Maryland, 2020-11-06), QTY30108 (USA/New York, 2021-03-30), QXS96145 (USA/Vermont, 2021-06-26), QQA02925 (USA/Maryland, 2020-11-11), QUA15581 (USA/New York, 2021-01-04), QUB14694 (USA/Michigan, 2021-03-24), QSJ39354 (USA/Florida, 2021-02-10), QWE91986 (USA/California, 2020-08-29), UEH28437 (USA/Colorado, 2020-09-16), UFD42885 (USA/Illinois, 2021-10-30); **f.** QUD12271 (USA/Ohio, 2021-03/27), QKI36860 (China/Guangzhou, 2020-02-21), QWT91543 (USA/New Mexico, 2020-11-10); UBY86923 (USA/Connecticut, 2021-09-08); QSX87276 (USA/Pennsylvania, 2021-02-25); **g.** UER91358 (USA/Maryland, 2021-10-22); QYF68609 (Bahrain, 2021-06-26); QXG22727 (Bahrain, 2021-06-29); QZP72779 (USA/Alabama, 2021-07-30); UEV0558 (USA/New York, 2021-10-24); UEH58452 (USA/North Carolina, 2021-10-01); **h.** UDN20252 (USA/Minnesota, 2021-10-15), UED44100 (USA/West Virginia, 2021-10-19), UFI73000 (USA/Minnesota, 2021-11-10), UBC64522 (USA/Wisconsin, 2021-07-21), UDL67371 (USA/Kentucky, 2021-09-29), QZS17582 (USA/Maryland, 2021-08-05), UDL43890 (USA/Georgia, 2021-09-26), UFM61198 (USA/Arizona, 2021-10-27), QZS20143 (USA/South Carolina, 2021-08-06), UCN28493 (USA/Minnesota, 2021-09-27), UBD47947 (USA/Virginia, 2021-09-05), UDA13813 (USA/Minnesota, 2021-10-04), QTU42716 (USA/Minnesota, 2021-01-12), UEM47659 (USA/Michigan, 2021-09-23), UDI49835 (USA/Tennessee, 2021-09-21), UDE93735 (USA/Idaho, 2021-10-09), UDA13516 (USA/Minnesota, 2021-10-03), UEQ92175 (USA/Indiana, 2021-10-25), QZS14557 (USA/Maryland, 2021-08-06), UDE67664 (USA/New Mexico, 2021-09-20), UFI75910 (USA/Minnesota, 2021-11-11), UAX90256 (USA/California, 2021-08-12), UBD49836 (USA/Tennessee, 2021-08-31), UDK33208 (USA/North Carolina, 2021-09-23), UDE66381 (USA/Missouri, 2021-09-20), QYS45691 (USA/Minnesota, 2021-07-28), UCK68269 (USA/Minnesota, 2021-08-30), QYY74618 (USA/Georgia, 2021-07-28), UBX72566 (USA/Maryland, 2021-09-01), UDE84787 (USA/South Carolina, 2021-09-21), UEU84720 (USA/Minnesota, 2021-10-31), UDK33636 (USA/Tennessee, 2021-09-23), QZT93722 (USA/Colorado, 2021-08-10); **i.** UDG72468 (USA/Tennessee, 2021-09-13), QTS66472 (USA/California, 2021-03-17), QYY75081 (USA/Georgia, 2021-07-28); QZI47484 (USA/California, 2021-08-16); QUA19334 (USA/Louisiana, 2021-02-02); UBY63352 (USA/Arizona, 2021-09-05); **j.** G (at position 53) in valine codon is replaced by U yielding an L: QTC62354 (USA/California, 2021-02-18); QUF36008 (USA/Michigan, 2021-03-31), QVV00321 (USA/Michigan, 2021-04-22); QSN9715 (USA/Pennsylvania, 2021-02-09); **k.** QQA02853 (USA/Maryland, 2021-11-11), QQS74355 (USA/Maryland, 2021-01-06), QQP31844 (USA/Maryland, 2020-12-29), QQA02817 (USA/Maryland, 2020-11-11); QYM26975 (USA/California, 2020-12-21); QTJ72925 (USA/Massachusetts, 2021-03-10); **l.** QUA15500 (USA/Kansas, 2021-01-02), QTM64537; QVM25420 (USA/Washington, 2021-02-10); QVM48721 (USA/Washington, 2021-03-17).

**Supplemental Figure to Figure 2 in main text**

**A.**

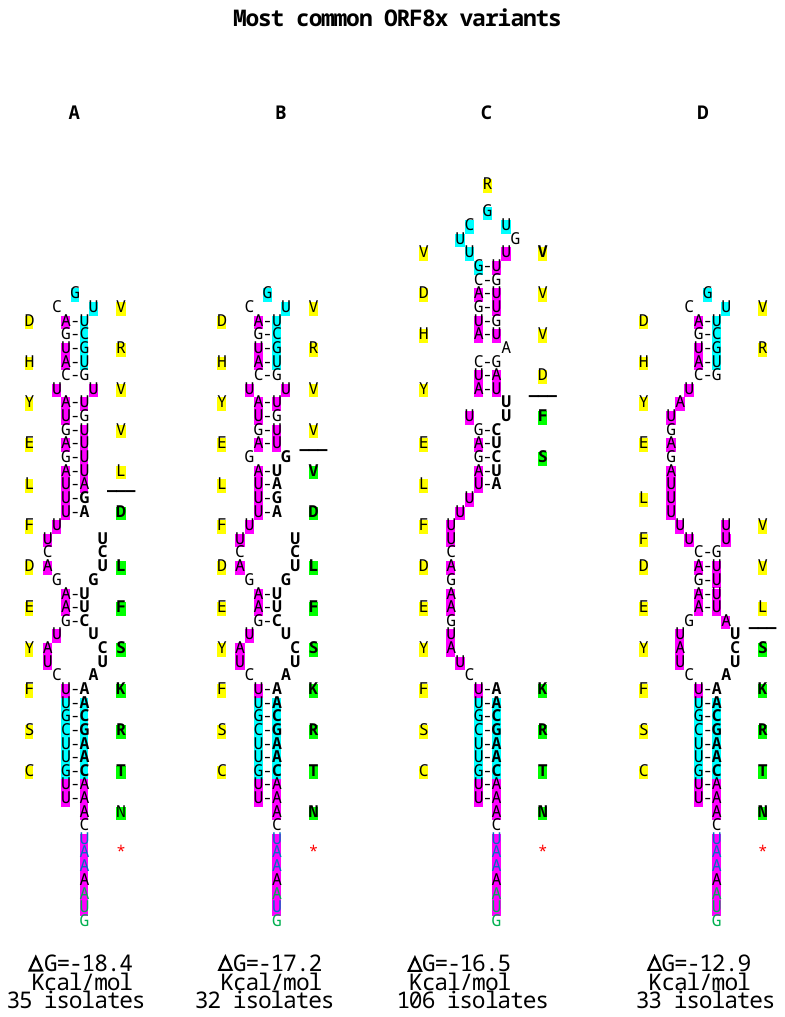

**B.**

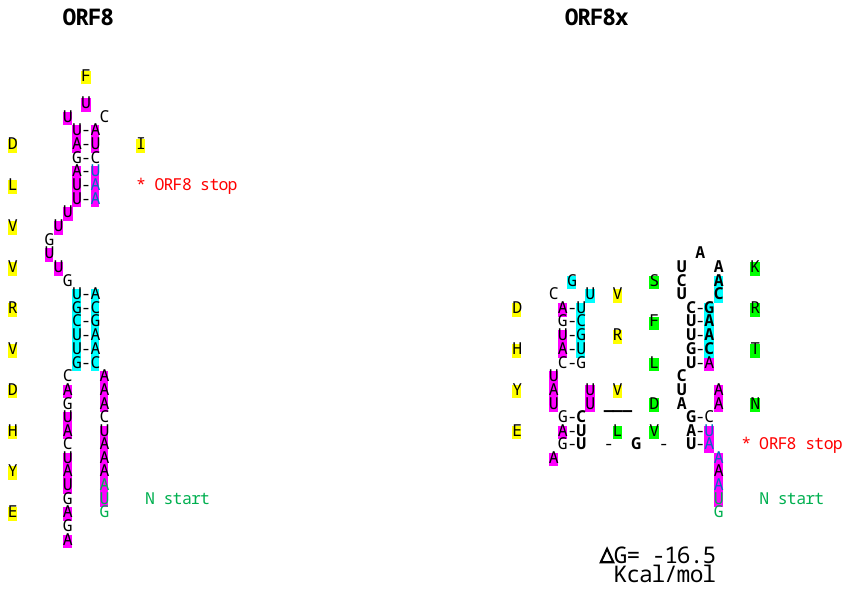

**Legend to Supplemental Figure to Figure 2 in main text**

**A. Most common ORF8 variants (ORF8x) with 5’-UTR-derived intragenomic rearrangement.**

Adenosine and uridine residues are highlighted in fuchsia, while TRS-B and cTRS-B sequences are highlighted in yellow. The nucleotide and amino acid sequence of the 5’-UTR-derived insertions are highlighted in green, while the amino acid sequence for the reference ORF8 is highlighted in yellow. Red asterisk denote stop codons. Number and distribution of collection sites for the most common ORF variants are: A. 35 isolates (all USA): California (5), Washington (3), Arizona (3), New York (3), Pennsylvania (2), Florida (2), Texas (2), West Virginia (2), Arkansas (1), South Carolina (1), North Carolina (1), Kansas (1), Georgia (1), Massachusetts (1), New Jersey (1), Nevada (1), Alabama (1), Minnesota (1), Illinois (1), Missouri (1). B. 33 isolates (all USA): Michigan (12), Florida (5), Virginia (2), California (2), Texas (2), Alabama (1), Arizona (1), Ohio (1), New Jersey (1), Illinois (1), Minnesota (1), Massachusetts (1), Arkansas (1), Kentucky (1), Pennsylvania (1). **C.** 106 isolates: California (14), Florida (9), Maryland (8), Pennsylvania (8), Colorado (5), Michigan (5), Minnesota (5), Texas (5), Arizona (3), North Carolina (3), South Carolina (3), Illinois (3), New Jersey (3), Virginia (3), New York (3), New Hampshire (2), Massachusetts (2), West Virginia (2), Tennessee (2), Georgia (2), Indiana (2), Alabama (1), Missouri (1), Utah (1), New Mexico (1), Maine (1), Vermont (1), Kansas (1), Wisconsin (1), District of Columbia (1), Kenya (1), Pakistan (1). C. 33 isolates (all US): Minnesota (10), Maryland (3), Tennessee (3), Georgia (2), South Carolina (2), West Virginia (1), Wisconsin (1), Kentucky (1), Arizona (1), Virginia (1), Michigan (1), Idaho (1), Indiana (1), New Mexico (1), California (1), North Carolina (1), Missouri (1), Colorado (1). **B.** **Alternative structure to that shown in Panel A with TRS-B binding to second cTRS-B.** The free minimum energy is similar to that for TRS-B binding to the first cTRS-B.

**Supplemental legend to Figure 4 in main text**

**SARS-CoV-2 variants (collection site and date in parentheses) with 5’-UTR-derived insertions modifying the SR region of the N protein; those shown in Figure 4 in main text are highlighted**

**a.** QTO33828 (EPI_ISL_1493720, USA/Texas, 2021/03/14; VOC Alpha GRY [B.1.1.7+Q.*] first detected in the UK), QWQ74880 (EPI_ISL_2528376, USA/Tennessee, 2021-06-03; VOC Gamma GR/501Y.V3 (P.1+P.1.*) first detected in Brazil/Japan), EPI_ISL_2252927 (Canada/Ontario, 2021-04; VOC Gamma GR/501Y.V3 (P.1+P.1.*) first detected in Brazil/Japan); **b.** EPI_ISL_3434731 (Brazil/Espirito Santo, 2021-07-18; VOC Gamma GR/501Y.V3 (P.1+P.1.*) first detected in Brazil/Japan), EPI_ISL_4104739 (Brazil/Sao Paulo, 2021-04-11; VOC Gamma GR/501Y.V3 (P.1+P.1.*) first detected in Brazil/Japan), EPI_ISL_2466134, Brazil/Rio Grande do Sul, 2021-04-15; VOC Gamma GR/501Y.V3 (P.1+P.1.*) first detected in Brazil/Japan), EPI_ISL_4276898 (USA/North Carolina, 2021-07-29; VOC Alpha GRY (B.1.1.7+Q.*) first detected in the UK), EPI_ISL_6050129 (USA/Florida, 2021-04-27; VOC Alpha GRY (B.1.1.7+Q.*) first detected in the UK), EPI_ISL_6050433 (USA/Florida, 2021-04-27; VOC Alpha GRY (B.1.1.7+Q.*) first detected in the UK), EPI_ISL_5676487 (Brazil/Sao Paulo, 2021-07-11; VOC Gamma GR/501Y.V3 (P.1+P.1.*) first detected in Brazil/Japan), EPI_ISL_3487143 (Canada/British Columbia, 2021-05-20; VOC Gamma GR/501Y.V3 (P.1+P.1.*) first detected in Brazil/Japan), EPI_ISL_5649672 (Brazil/Sao Paulo, 2021-05-23; VOC Gamma GR/501Y.V3 (P.1+P.1.*) first detected in Brazil/Japan), EPI_ISL_2493472 (Brazil/Sao Paulo, 2021-05-16; VOC Gamma GR/501Y.V3 (P.1+P.1.*) first detected in Brazil/Japan), EPI_ISL_5802386 (Brazil/Sao Paulo, 2021-04-11; VOC Gamma GR/501Y.V3 (P.1+P.1.*) first detected in Brazil/Japan), EPI_ISL_5802523, Brazil/Sao Paulo, 2021-04-20; VOC Gamma GR/501Y.V3 (P.1+P.1.*) first detected in Brazil/Japan), EPI_ISL_5669813 (Brazil/Sao Paulo, 2021-07-05; VOC Gamma GR/501Y.V3 (P.1+P.1.*) first detected in Brazil/Japan), EPI_ISL_5663882 (Brazil/Sao Paulo, 2021-06-30; VOC Gamma GR/501Y.V3 (P.1+P.1.*) first detected in Brazil/Japan), EPI_ISL_5651350 (Brazil/Sao Paulo, 2021-07-17; VOC Gamma GR/501Y.V3 (P.1+P.1.*) first detected in Brazil/Japan), EPI_ISL_5649266 (Brazil/Sao Paulo, 2021-06-20; VOC Gamma GR/501Y.V3 (P.1+P.1.*) first detected in Brazil/Japan), EPI_ISL_5647661 (Brazil/Sao Paulo, 2021-06-13; VOC Gamma GR/501Y.V3 (P.1+P.1.*), EPI_ISL_7615326 (Brazil/Tocantis, 2021-08-22; VOC Gamma GR/501Y.V3 (P.1+P.1.*) first detected in Brazil/Japan), EPI_ISL_3987895 (Chile/Ñuble, 2021-08-10; VOC Gamma GR/501Y.V3 (P.1+P.1.*) first detected in Brazil/Japan), EPI_ISL_3987894 (Chile/Ñuble, 2021-08-10; VOC Gamma GR/501Y.V3 (P.1+P.1.*) first detected in Brazil/Japan), EPI_ISL_4417129 (Peru/Lima, 2021-08-11; VOC Gamma GR/501Y.V3 (P.1+P.1.*) first detected in Brazil/Japan), EPI_ISL_4414506 (Brazil/Sao Paulo, 2021-07-02; VOC Gamma GR/501Y.V3 (P.1+P.1.*) first detected in Brazil/Japan), EPI_ISL_4413570 (Brazil/Piaui, 2021-08-10; VOC Gamma GR/501Y.V3 (P.1+P.1.*) first detected in Brazil/Japan), EPI_ISL_3259363 (Brazil/Goias, 2021-07-12; VOC Gamma GR/501Y.V3 (P.1+P.1.*) first detected in Brazil/Japan), EPI_ISL_3049003 (Brazil/Sao Paulo, 2021-06-15; VOC Gamma GR/501Y.V3 (P.1+P.1.*) first detected in Brazil/Japan), EPI_ISL_3832398 (Brazil/Rio de Janeiro, 2021-03-15; VOC Gamma GR/501Y.V3 (P.1+P.1.*) first detected in Brazil/Japan), EPI_ISL_2345318 (Brazil/Sao Paulo, 2021-03-21; VOC Gamma GR/501Y.V3 (P.1+P.1.*) first detected in Brazil/Japan), EPI_ISL_2345318 (Brazil/Sao Paulo, 2021-03-21; VOC Gamma GR/501Y.V3 (P.1+P.1.*) first detected in Brazil/Japan), EPI_ISL_3664170 (Brazil/Tocantins, 2021-05-10; VOC Gamma GR/501Y.V3 (P.1+P.1.*) first detected in Brazil/Japan), EPI_ISL_4347353 (USA/Florida, 2021-05-08; VOC Alpha GRY (B.1.1.7+Q.*) first detected in the UK), EPI_ISL_1916295 (India/Maharashtra, 2021-01-06; B.1.1 (Pango v.3.1.20 2022-02-02)), EPI_ISL_1795118 (Brazil/Sao Paulo, 2021-01-06; VOC Gamma GR/501Y.V3 (P.1+P.1.*) first detected in Brazil/Japan), EPI_ISL_295866 (USA/Michigan, 2021-04-16), EPI_ISL_1320623 (USA/North Carolina, 2021-03-02; B.1.1.519 (Pango v.3.1.20 2022-02-02)).

**Supplemental legend to Figure 5 in main text**

**SARS-CoV-2 variants (collection site and date in parentheses) with 5’-UTR-derived insertions modifying the N-terminal NiRAN domain of the RNA-dependent RNA polymerase (nsp12); those shown in Figure 5 in main text are highlighted**

**a.** QVL75820 (EPI_ISL_1209225, USA/Washington, 2021-03-28), EPI_ISL_1524008 (USA/Washington, 2021-03-28) **b.** UHP90975 (USA/Wisconsin, 2021-12-13), UCX37945 (USA/Wisconsin, 2021-09-20), UET58811 (USA/North Carolina, 2021-10-22), QSF06151 (USA/California, 2021-02-09), QVJ61220 (USA/Tennessee, 2021-04-13), QRW84488 (USA/Massachusetts, 2021-01-17), UBA27920 (USA/Connecticut, 2021-08-26), QTQ63154 (USA/Oregon, 2021-03-05), QUA74029 (USA/Tennessee, 2021-03-30), UHF32869 (USA, 2021-10-07), QSF02035 (USA/North Carolina, 2021-02-04), UCU50446 (USA, 2021-08-07), UDI24491 (USA/Michigan, 2021-09-16),UCX29694 (USA/Wisconsin, 2021-09-17), QUB30213 (USA/Tennessee, 2021-03-16), QTP86894 (USA/Tennessee, 2021-03-23), UHT32237 (USA/Texas, 2021-12-15), UDN70797 (USA/New Jersey 2021-10-02), QSG76884 (USA/Louisiana, 2021-02-11), UHQ26009 (USA/West Virginia, 2021-1206), QQK89755 (USA/Texas, 2020-06-20), QTY89177 (USA/Tennessee, 2021-03-18), UCU60974 (Kenya, 2020-11-12), QQK89671 (USA/Texas, 2020-06-20), UDQ13548 (USA/Massachusetts, 2021-09-30), UHH55298 (USA/Colorado, 2021-10-07), UFL45530 (USA/Wisconsin, 2021-11-07), QUF14763 (USA/Tennessee, 2021-04-09), UIB16394 (USA/California, 2021-12-23), QSN79388 (USA/Louisiana, 2021-02-10), UEL74880 (USA/New York, 2021-10-30), QTC17106 (USA/Pennsylvania, 2021-02-27), QZU12979 (USA/Maryland, 2021-08-08), QTP87948 (USA/Tennessee, 2021-03-23), QXH28738 (USA/Alabama, 2021-04-16), QTQ45244 (USA/Massachusetts, 2021-03-02), UCK93316 (Mexico, 2021-02-10), UDN75390 (USA/Wisconsin, 2021-10-14), UDI22517 (USA/Michigan, 2021-09-16), UEH77441 (USA/Idaho, 2021-10-06), UEI88362 (USA/New York, 2021-10-22), UFO07215 (China/Guangzhou, 2021-05-26), QYR09303 (USA/California, 2021-07-22), QTG30874 (USA/Pennsylvania, 2021-03-03), UFB48090 (USA/Ohio, 2021-11-01), UEN28867 (USA/North Carolina, 2021-10-10), UCX35525 (USA/Michigan, 2021-09-19), UDI24219 (USA/Michigan, 2021-09-16), UFK35880 (USA/Colorado, 2021-10-27), QVP10190 (USA/Tennessee, 2021-03-09), QQD89619 (USA/Texas, 2020-06-19), UDL68518 (USA/Utah, 2021-09-30), UFS19776 (USA/Colorado, 2021-10-19), UFC62618 (USA/North Carolina, 2021-10-17), UFD31246 (USA/Wisconsin, 2021-11-03), UIT57972 (USA/California, 2021-02-15), UBF49486 (USA/Florida, 2021-08-17), QTS23884 (USA/Pennsylvania, 2021-03-17), UFM23938 (USA/Wisconsin, 21021-11-12), QWT32066 (USA/Florida, 2021-04-13), UCL05955 (Mexico, 2021-02-10), QWT32449 (USA/Florida, 2021-04-20), UHY96156 (USA/Indiana, 2021-12-20), QSO02183 (USA/Georgia, 2021-02-15), UHA91620 (USA/Michigan, 2021-02-27), QQD89259 (USA/Texas, 2020-06-19), UFB17148 (USA/California, 2021-09-18), UBI86079 (USA/Washington, 2021-08-24), UFC48585 (USA/Wisconsin, 2021-10-15), QUB35876 (USA/Tennessee, 201-03-18), UET19478 (USA/California, 2021-10-22), QTY89153 (USA/Tennessee, 2021-03-18), QTZ12792 (USA/Tennessee, 2021-03-25), UDN76025 (USA/Wisconsin, 2021-10-15), QUP24690 (USA/Tennessee, 2021-04-03), QRW75822 (USA/Georgia, 2021-01-19), QZQ18874 (USA/Maryland, 2021-07-22), UEH80397 (USA/New Jersey, 2021-10-04), QRW35316 (USA/California, 2021-02-08), QTD08611 (USA/Pennsylvania, 2021-03-01), QZN42224 (USA/Massachusetts, 2021-08-19), QTH26806 (USA/Texas, 2021-03-15), UBA30158 (USA/Connecticut, 2021-08-24), UEH42753 (USA/Utah, 2021-09-29), QRW35868 (USA/California, 2021-02-07), UHN79885 (USA/Wisconsin, 2021-12-09), QSN82400 (USA/Utah, 2021-02-09), UEK36591 (USA/California, 2021-08-25), UGO85135 (USA/Michigan, 2021-03-26), QZN59551 (USA/Massachusetts, 2021-08-21), QTP89015 (USA/New Mexico, 2021-03-22), UHU01313 (USA/New Jersey, 2021-12-09), QUB09583 (USA/Tennessee, 2021-03-22), QUA79262 (USA/Indiana, 2021-03-30), QZR27774 (USA/New Jersey, 2021-08-08), QUF39210 (USA/Virginia, 2021-03-30), QTK00317 (USA/Tennessee, 2021-03-09), QQY92455 (USA/Massachusetts, 2020-10-30), QTP82254 (USA/Indiana, 2021-03-18), QUC73664 (USA/New Jersey, 2021-03-26), UDQ12989 (USA/Texas, 2021-09-30), QSN95378 (USA/Massachusetts, 201-02-11), UHV84541 (USA/South Dakota, 2021-12-17), QTN63096 (USA/Tennessee, 2021-03-16), UEN25043 (USA/California, 2021-10-09), UDI27036 (USA/Michigan, 2021-09-17), UAU82483 (USA/Mississippi, 2021-08-23), UCK93340 (Mexico, 2021-02-12), QSN81011 (USA/Massachusetts, 2021-02-09), QWE98942 (USA/Tennessee, 201-04-01), QUP23047 (USA/Pennsylvania, 2021-04-06), UCK93376 (Mexico, 2021-02-12), QUE01559 (USA/Michigan, 2021-03-25), UIG95057 (USA/Texas, 2020-07-11), UCK93328 (Mexico, 2021-02-12), QSL68204 (USA/Massachusetts, 2021-02-15), UBJ41712 (EPI_ISL_4210459, USA/Colorado, 2021-07-17), UCN32677 (USA/Wisconsin, 2021-09-07), UHB31750 (USA/Wisconsin, 2021-12-06), UGW49737 (USA/Rhode Island, 2021-12-02), QTC69545 (USA/North Carolina, 2021-02-19), QSE23148 (USA/Nevada, 2021-02-02), QTX67843 (USA/Pennsylvania, 2021-02-28), UEO69019 (USA/Vermont, 2021-11-02), QTI92251 (USA/Texas, 2021-03-11), UED49901 (USA/Alabama, 2021-10-11), QUS65424 (USA/Michigan, 2021-04-15), UCL09275 (Mexico, 2021-02-08), UCU26113 (USA/Colorado, 2021-09-15), UBE28012 (USA/Tennessee, 2021-08-31), QZM41370 (USA/Maryland, 2021-08-02), QWP91207 (USA/Utah, 2021-01-12), UCK93017 (Mexico, 2021-02-11), UFL58409 (USA/Georgia, 2021-11-09), UAQ08726 (USA/Kentucky, 2021-08-19), QVQ47721 (USA/Utah, 2021-02-17), UGO30951 (USA/Michigan, 2021-04-09), UCK93268 (Mexico, 2021-02-11), QUC95975 (USA/Wisconsin, 2021-03-29), UCK93256 (Mexico, 2021-02-11), QST20745 (USA/California, 2021-02-16), UFR30090 (USA/Texas, 2021-11-13), UFY60772 (USA/Wisconsin, 2021-11-16), UCK93208 (Mexico, 2021-02-11), UCK93412 (Mexico, 2021-02-09), QYZ51361 (USA/California, 2021-07-22), QSY38005 (USA/Utah, 2021-02-11), QRX36206 (USA/Georgia, 2021-01-26), UCU51547 (USA/California, 2021-08-05), UAJ86375 (USA/Massachusetts, 2021-08-28), UCU60808, UDE70831, UEJ48203, UFG29076, UDN70720, QTS15625, UCV75209, QOL76451, QTP86918, UBI85816 (USA/Washington, 2021-08-25), UHU25439 (USA/Colorado, 2021-12-15), QZH95271 (USA/California, 2021-08-14), UCA83849 (USA/Tennessee, 2021-08-25), UAU14550 (USA/California, 2021-08-07), QTM59778 (USA/California, 2021-03-10), UAB04361 (USA/California, 2021-08-24), QZO45287 (USA/California, 2021-08-16), QZG94711 (USA/California, 2021-08-09), QTM28698 (USA/California, 2021-03-14), UAA69987 (USA/California, 2021-08-20), QZC68055 (USA/California, 2021-08-06), QTM60426 (USA/California, 2021-03-10), QTM35574 (USA/California, 2021-03-19), QZI23057 (USA/California, 2021-08-11), UAC34418 (USA, California, 2021-08-19), QZS28746 (USA/California, 2021-08-11), QZI43852 (USA/California, 2021-08-16), QTM53706 (USA/California, 2021-03-11), QZH97195 (USA/California, 2021-08-14), QTM62250 (USA/California, 2021-03-10), QZS26314 (USA/California, 2021-08-11), QTM46434 (USA/California, 2021-03-09), QZH93082 (USA/California, 2021-08-13), QTM62430 (USA/California, 2021-03-10), QTM28638 (USA/California, 2021-03-14), QTM61734 (USA/California, 2021-03-11), QTM55686 (USA/California, 2021-03-21), QTM51006 (USA/California, 2021-03-03), QZG92437 (USA/California, 2021-08-09), QTQ57981 (USA/California, 2021-03-25), QZC67661 (USA/California, 2021-08-05), UAU22498 (USA/California, 2021-08-09), UAC24682 (USA/California, 2021-08-18), UAU15783 (USA/California, 2021-08-07), QYO40026 (USA/California, 2021-07-26), UAM59368 (USA/California, 2021-08-23), UEW99265 (USA/Minnesota, 2021-09-07), QTM59310 (USA/California, 2021-03-04), QTW87379 (USA/California, 2021-04-05), QTM53802 (USA/California, 2021-03-11), UCM07937 (USA/New Jersey, 2021-09-24), UIA74851 (USA/California, 2021-12-11), QWS07135 (USA/Massachusetts, 2021-05-26), QVO70308 (USA/Massachusetts, 2021-05-10), UIB88512 (USA/California, 2021-12-08), UHD98968 (USA/Washington, 2021-12-10), UHS95021 (USA/California, 2021-12-04), QZI30998 (USA/California, 2021-08-12), QWE74279 (USA/Rhode Island, 2021-05-20), QWT11941 (USA/Rhode Island, 2021-05-19), UHS99543 (USA/California, 2021-12-06), QVU19426 (USA/Tennessee, 2021-05-14), QVK91899 (USA/California, 2021-04-29), UIC02954 (USA/California, 2021-12-11), UDW21718 (USA/Florida, 2021-06-30), UGZ85477 (USA/Washington, 2021-12-02), QVV52552 (USA/Pennsylvania, 2021-05-06), QVU40092 (USA/Tennessee, 2021-05-14), UGV83346 (USA/California, 2021-11-22), QVE27167 (USA/Minnesota, 2021-04-08), UIG78628 (USA/New York, 2021-12-20), UHZ77630 (USA/New York, 2021-12-23), UIO66406 (USA/Colorado, 2021-12-22), UEP72746 (USA/Utah, 2021-10-06), QTQ57299 (USA/California, 2021-03-25), QTW87367 (USA/California, 2021-04-05), QXI81186 (USA/Utah, 2021-01-20), UEZ67112 (USA/Florida, 2021-08-21), QTM59923 (USA/California, 2021-03-10), UAB04274 (USA/California, 2021-08-24), QTM50971 (USA/California, 2021-03-03), QVV13623 (USA/Tennessee, 2021-04-22), UIS90595 (USA/California, 2021-02-23), QSH79120 (USA/Massachusetts, 2020-12-23), QSH79072 (USA/Massachusetts, 2020-12-22), UEO69832 (USA/Vermont, 2021-11-02), QTQ43251 (USA/Massachusetts, 2021-02-16), QTY91588 (USA/Virginia, 2021-03-18), UFD88645 (USA/Florida, 2021-09-01). **c.** UAQ66644 (USA/Virginia, 2021-07-21), QWO72252 (USA/Utah, 2021-01-16), UDN79241 (USA/California, 2021-10-17), UFB02276 (USA/California, 2021-09-28), UBR04551 (EPI_ISL_1700692, USA/California, 2021-04-01), QOT58454 (Australia/Victoria, 2020-08-06), QNO32118 (USA/FL, 2020-07-10), QTF73897 (USA/Washington, 2020-04-17), QOT52503 (Australia/Victoria, 2020-07-29), QOT65712 (Australia/Victoria, 2020-08-04), UAL05318 (USA/New Jersey, 2021-08-04), QVL90312 (USA/Alaska, 2021-02-15), QUG27104 (USA/California, 2021-04-04), QSH79834 (USA/Massachusetts, 2021-02-05), QOT55131 (Australia/Victoria, 2020-08-11), QNP04397 (Australia/Victoria, 2020-07-07), QVJ30557 (USA/Ohio, 2021-04-10), QKV37606 (Australia/Victoria, 2020-03-23), UDN80965 (USA/California, 2021-10-18), UBU58048 (USA/Vermont, 2021-09-24), QTP78529 (USA/Ohio, 2021-03-24), QNO31650 (USA/Florida, 2020-06-03), QIZ13153 (USA/Washington, 2020-03-23), QTP21880 (USA/Pennsylvania, 2021-03-08), QZM66495 (USA/North Carolina, 2021-08-05), UHP32610 (USA/Colorado, 2021-11-26), QOQ08762 (Australia/Victoria, 2020-08-14), QUP30365 (USA/North Carolina, 2021-04-02), UGV53892 (EPI_ISL_7332674, USA/Colorado, 2021-11-18), QKV39238 (USA/Washington, 2020-04-28), QPN00245 (USA/Virginia, 2020-10), QRW69490 (USA/Kansas, 2021-01-02), UFT01738 (EPI_ISL_6911013, USA/Colorado, 2021-11-07), UBN73346 (USA/California, 2021-07-21), QTW98676 (USA/Ohio, 2021-04-08), QZM69710 (USA/North Carolina, 2021-08-07), QOT67656 (Australia/Victoria, 2020-08-11), QTY90842 (USA/Ohio, 2021-03-25), QZJ82123 (USA/Arizona, 2020-11-05), QQW60892 (EPI_ISL_862697, USA/New Mexico, 2021-01-11), QSM36221 (USA/Virginia, 2021-02), QSV97066 (USA/Georgia, 2021-02-25), UHJ22250 (USA/Washington, 2021-11-29), QYB96896 (USA/Nevada, 2021-07-06), QSH74103 (USA/Washington, 2021-02-02), UBG29595 (USA/Pennsylvania, 2021-03-08), UCP53771 (USA/Arizona, 2020-10-29), UFD69116 (USA/Nebraska, 2021-11-01), UET89315 (USA/New Mexico, 2021-10-20), QXL30633 (USA/Nevada, 2021-06-27), QNP05333 (Australia/Victoria, 2020-07-22), UAJ51444 (EPI_ISL_4000736, USA/Texas, 2021-08-07), UIA22551 (USA/Illinois, 2021-12-16), UCA85430 (USA/New Jersey, 2021-08-24), UDA94630 (USA/California, 2021-01-14), QUF14319 (USA/California, 2021-03-30), UEI47887 (USA/Texas, 2021-10-18), UDI32583 (USA/California, 2021-09-23), UEX06825 (USA/Minnesota, 2021-09-15), UEX08512 (USA/Minnesota, 2021-09-15), UDB02101 (Mexico/Baja California, 2020-05-12), UAU26527 (USA/Washington, 2021-07-30), UEF01099 (USA/Vermont, 2021-10-27), UFL76189 (USA/Kentucky, 2021-11-10), UBR04479 (EPI_ISL_1677791, USA/California, 2021-03-29), QTC82985 (USA/Georgia, 2021,02-21), UBS20118 (USA/Washington, 2021-09-06), QUP33452 (USA/Michigan, 2021-04-07), UGY44799 (USA/North Carolina, 2021-08-21), UBR04503 (USA/California, 2021-03-29), QSL69846 (USA/California, 2021-02-08), UCB64272 (USA/North Carolina, 2021-08-25), QTC77717 (USA/North Carolina, 2021-02-21), UCQ05117 (USA/Colorado, 2021-09-13), UBR04539 (EPI_ISL_1700691, USA/California, 2021-0401), QSN97897 (USA/Ohio, 2021-02-04), QUG29808 (USA/Ohio, 2021-04-07), QSX71723 (USA/South Carolina, 2021-01-22), QOT47919 (Australia/Victoria, 2020-08-11), UGW00470 (USA/Vermont, 2021-12-01), UBG86513 (USA/Texas, 2021-06-15), QOT61081 (Australia/Victoria, 2020-08-11), QNP05765 (Australia/Victoria, 2020-07-14), UDO75557 (USA/California, 2021-09-02), QZJ99124 (USA/Arizona, 2021-11-06), QYC02481 (USA/Virginia, 2021-07), QSQ87317 (Spain, 2021-02-06), QSO34191 (USA/Washington, 2021-02-01), UCL05620 (Mexico, 2021-02-10), UDH85977 (USA/Michigan, 2021-10-09), QZM70941 (USA/New York, 2021-08-05), UHQ63083 (USA/Colorado, 2021-11-26), QNP05369 (Australia/Victoria, 2020-07-21), UBT01574 (USA/Vermont, 2021-09-20), QXT01211 (USA/Nevada, 2021-07-03), UGN24690 (USA/Kentucky, 2021-09-08), QZK03982 (USA/Arizona, 2021-11-13), UET58952 (USA/California, 2021-10-22), UEC53187 (USA/California, 2021-10-19), UFN90271 (USA/Vermont, 2021-11-16), UBT66370 (USA/Rhode Island, 2021-09-13), QSX86679 (USA/Connecticut, 2021-02-25), UFY47349 (USA/California, 2021-11-15), QSO06088 (USA/Ohio, 2021-02-10), UEV59764 (USA/Colorado, 2021-10-14), UHR78108 (USA/Arizona, 2021-12-07), QZZ79447 (USA/Minnesota, 2021-08-01), QKV37366 (Australia/Victoria, 2020-03-24), UBC57025 (USA/Oregon, 2021-08-02), QSV31123 (USA/Ohio, 2021-02-03), UFH09963 (USA/California, 2021-11-10), QYK94763 (USA/Arkansas, 2021-07-21), QZR23215 (USA/New Jersey, 2021-08-05), QVJ29898 (USA/California, 2021-04-09), QOQ08846 (Australia/Victoria, 2020-08-15), UFC64772 (USA/Georgia, 2021-10-17), UDK29334 (USA/Oklahoma, 2021-09-22), QNP05189 (Australia/Victoria, 2020-07-22), QRN62752 (USA/Washington, 2020-04-17), QYN93822 (USA/Minnesota, 2021-03-17), UIT15841 (USA/California, 2021-05-07), UFM46861 (USA/Texas, 2021-11-07), UDK29092 (USA/Georgia, 2021-09-22), QUX40238 (USA/Ohio, 2021-04-19), QOT56895 (Australia/Victoria, 2020-08-09), QZK76656 (USA/Arizona, 2021-10-27), UDK82950 (USA/Georgia, 2021-09-20), UFY09470 (USA/Colorado, 2021-11-08), UBR04134 (EPI_ISL_1664598, USA/California, 2021-03-16), QNA41452 (USA/Massachusetts, 2020-05-05), QTG62009 (USA/Minnesota, 2021-03-11), UCI03387 (USA/California, 2021-08-23), UFD69226 (USA/Michigan, 2021-11-01), UHL52806 (USA/Massachusetts, 2021-12-11), QZU53928 (USA/Mississippi, 2021-08-02), UFH62441 (USA/Washington, 2021-11-08), QWO67801 (USA/Texas, 2021-05-25), QOT66936 (Australia/Victoria, 2020-08-04), UBG22777 (USA/Minnesota, 2021-08-12), UAZ00628 (USA/Colorado, 2021-08-10), QKV37702 (Australia/Victoria, 2020-03-25), QOT49983 (Australia/Victoria, 2020-08-18), UAK10370 (USA/Massachusetts, 2021-09-01), QRW53158 (USA/unknown, 2021-12-128), UGA98207 (USA/Pennsylvania, 2021-11-22), QNP05621 (Australia/Victoria, 2020-07-23), QYV26986 (USA/Ohio, 2021-02-19), QWQ56523 (Egypt, 2021-05-03), UCO32484 (USA/South Carolina, 2021-09-08), UHY49505 (USA/California, 2021-11-21), QWQ09843 (USA/New Mexico, 2021-03-03), UBE59357 (USA/Nevada, 2021-06-05), QYP28652 (USA/Florida, 201-07-16), UFP72527 (USA/California, 2021-11-02), QOQ01576 (Australia/Victoria, 2020-07-22), UFB68519 (USA/Washington, 2021-11-04), QZQ30539 (USA/Utah, 2021-08-19), UFD04864 (USA/Maryland, 2021-10-20), UFW04312 (USA/District of Columbia, 2021-11-18), UFB50646 (USA/New Hampshire, 2021-11-01), QSN92962 (USA/Ohio, 2021-02-11), QNP04829 (Australia/Victoria, 2020-07-22), UEZ48750 (USA/Georgia, 2021-08-16), UHF44058 (USA/California, 2021-11-22), UAX63475 (USA/Connecticut, 2021-09-06), QWP94410 (USA/California, 2021-04-19), UHE84670 (USA/Connecticut, 2021-11-10), UFO57784 (USA/Colorado, 2021/10/30), QSO06053 (USA/Ohio, 2021-02-09), QTP21832 (USA/Pennsylvania, 2021-03-08), UIW34244 (USA/Idaho, 2021-10-07), UCL68725 (USA/Vermont, 2021-09-29), QNO72753 (Australia/Victoria, 2020-07-14), UFR51422 (USA/Maryland, 2021-11-13), UGP26607 (USA/California, 2021-11-16), UHJ22059 (USA/Pennsylvania, 2021-11-25), UHA50630 (USA/Massachusetts, 2021-12-03), UET95602 (USA/Nevada, 2021-10-23), QNP04469 (Australia/Victoria, 2020-07-11), UCP63823 (USA/Arizona, 2021-07-21), UDH04229 (USA/Utah, 2021-09-06), UID27151 (USA/Colorado, 2021-12-07), QKV37330 (Australia/Victoria, 2020-03-22), QNP04709 (Australia/Victoria, 2020-07-22), QVK75504 (USA/California, 2021-05-05), UAZ99719 (USA/New York, 2021-04-16), QZI50984 (USA/California, 2021-08-14), QYT26085 (USA/New York, 2021-07-27), UFY76065 (USA/California, 2021-11-17), UBZ72278 (USA/Minnesota, 2021-09-24), UAT65018 (USA/Texas, 2021-08-23), UBZ32334 (USA/Colorado, 2021-09-23), QUX63045 (USA/California, 2021-04-29), QTP21676 (USA/Pennsylvania, 2021-03-08), QUX63246 (USA/California, 2021-04-29), QWQ74846 (USA/Tennessee, 2021-06-03), UHE06838 (Kenya, 2021-04-23), QVM69527 (USA/Washington, 2021-04-10), QTP21868 (USA/Philadelphia, 2021-03-08), QUX78895 (USA/Philadelphia, 2021-04-01), QVO98143 (USA/Utah, 2021-03-09), QTP21928 (USA/Philadelphia, 2021-03-08), UHE10712 (USA/Kansas, 2021-11-19), UCS65869 (Kenya, 2021-04-27), QWQ63966 (USA/Massachusetts, 2021-05-28), QVH96002 (USA/Florida, 2021-04-24), QTT38102 (Austria, 2021-03-14), QVV41697 (USA/Texas, 2021-05-09), UAZ99443 (USA/California, 2021-05-06), UBG20578 (USA/California, 2021-07-07), QXR80990 (USA/New Jersey, 2021-03-15), QUO78350 (USA/North Carolina, 2021-04-15), QTW94838 (USA/Philadelphia, 2021-03-15), QUX78967 (USA/Philadelphia, 2021-04-01), QUX78919 (USA/Philadelphia, 2021-04-01), QUX78955 (USA/Philadelphia, 2021-04-01), QVU55471 (USA/Philadelphia, 2021-05-03), QWS10103 (USA/Connecticut, 2021-05-25), QUD29701 (USA/Pennsylvania, 2021-03-30), QTP21892 (USA/Philadelphia, 2021-03-08), UGW59478 (USA/California, 2020-12-22), UEI21061 (USA/California, 2020-10-27), QZK50146 (USA/Arizona, 2020-10-09), UHF53553 (USA/Illinois, 2021-11-22), UEC91489 (USA/Florida, 2021-10-04), UDD68618 (EPI_ISL_5259201, USA/Colorado, 2021-09-22), UBT74402 (EPI_ISL_4574730, USA/Colorado, 2021-08-09), QVV22431 (EPI_ISL_2229936, USA/Florida, 2021-03-05), UFO92620 (USA/Massachusetts, 2021-10-22), UIT75188 (USA/California, 2021-02-11), QUX63141 (USA/California, 2021-04-29), QUX74660 (USA/California, 2021-04-30), UCN77125 (USA/North Carolina, 2021-09-16), UGP27130 (USA/California, 2021-11-16), UBA22712 (USA/Florida, 2021-08-27), QXN03610 (USA/Nevada, 2021-06-24), QQW26461 (USA/Texas, 2020-10-04). **d.** UHS40780 (USA/California, 2021-09-23), UGO44472, UIW49833, UHZ07618, UHV76469; **e.** UFT72204 (USA/Colorado, 2021-10-27), EPI_ISL_1384819 (India/Maharashtra, 2021-02-12), EPI_ISL_1703925 (India/Maharashtra, 2021-02-07); **f.** QZM71485 (USA/New York, 2021-08-05), QTG28282 (USA/Pennsylvania, 2021-03-06), QVJ64469 (USA/California, 2021-04-14), QRX48545 (USA/Arizona, 2021-01-29), QTJ73204 (USA/Pennsylvania, 2021-03-11), QZX55999 (USA/Utah, 2021/01/28), QUH41414 (USA/California, 2021-04-06), UDA96392 (USA/California, 2021-01-19), QUE38893 (USA/California, 2021-03-31), QOW96201 (USA/Virginia, 2020-08), UDA78757 (USA/California, 2020-12-25), UBD95955 (USA/North Carolina, 2021-08-30), QXR83245 (USA/Pennsylvania, 2021-03-28), QTD04155 (USA/California, 2021-02-24), UDA84951 (Mexico/Baja California, 2020-12-28), UGY48417 (USA/North Carolina, 2021-08-31), UIH05096 (USA/Virginia, 2021-12-06), QVJ64457 (USA/California, 2021-04-14), QTM33114 (USA/California, 2021-03-15), QTM33066 (USA/California, 2021-03-15), QTM51534 (USA/California, 2021-03-02), QTM95171 (USA/California, 2021-03-03), QTM61710 (USA/California, 2021-03-11), QTM32886 (USA/California, 2021-03-15), QWS15650 (USA/Illinois, 2021-05-18), QVN36411 (USA/Illinois, 2021-05-09), QYV41469 (USA/New York, 2021-06-25), QVF50260 (USA/Illinois, 2021-04-26), QWB63840 (USA/Illinois, 2021-05-18), QTM38515 (USA/California, 2021-03-04), QXM96995 (USA/Rhode Island, 2021-06-22), QSN92016 (USA/New Hampshire, 2021-02-10); **g.** UBL67135 (USA/Maryland, 2021-09-07), UBL84726 (USA/Maryland, 2021-09-10), UDK37409 (USA/Maryland, 2021-09-21), QWJ83632 (USA/Florida, 2021-05-11); **h.** UBD35057 (USA/Texas, 2021-09-01), UHR18165 (USA/Maine, 2021-12-10), UBT64155 (USA/Massachusetts, 2021-09-20), UHF10135 (USA/California, 2021-11-15), QTJ77540 (USA/California, 2020-04-27), UBA73421 (USA/Maryland, 2021-09-02), QTJ82390 (USA/California, 2020-04-18), UDF93258 (USA/Maryland, 2021-10-04), UCB61890 (USA/Maryland, 2021-08-27), UBN88464 (USA/Maryland, 2021-04-10), QUI07616 (USA/New Jersey, 2021-04-08).

**Supplemental listing of coronaviruses without intragenomic rearrangements**

Using 5’ UTRs from reference isolates (in parentheses) as query sequences, no 5’-UTR insertions were detected in the genome bodies of other CoVs infecting humans including the Sarbecovirus β-CoV SARS-CoV-1 (NC_004718) and the human α-CoVs hCoV-229E (NC_002645, MW532103 and KU291448, subgenus Duvinacovirus ) and hCoV-NL63 (NC_005831, 686 isolates, subgenus Setracovirus). In addition, no insertions were found in: α-CoVs subgenus Tegacovirus feline CoV and infectious peritonitis virus (FECV and FIPV; NC_002306), transmissible gastroenteritis virus (TGEV; DQ811788, NC_038861: 151 isolates), Swine enteric coronavirus strain Italy/213306/2009 (NC_028806, 151 isolates), Canine coronavirus strain CCoV/NTU336/F/2008 (GQ477367, 151 isolates); Alphacoronavirus 1 strain 23/03 (KP849472, 151 isolates), Feline coronavirus strain FCoV/NTU156/P/2007 (GQ152141, 151 isolates), Feline coronavirus strain DF-2 (DQ286389, 151 isolates); Canine coronavirus strain (KC175339, 171 isolates), Feline coronavirus strain FCoV C1Je (DQ848678, 151 isolates), Feline coronavirus isolate 27C (KP143507), Feline coronavirus UU21 HQ012369), Feline coronavirus UU16 (FJ938058), Feline coronavirus UU24 (HQ012370), Feline coronavirus isolate Black (EU186072), subgenus Rhinacovirus severe acuate diarrhea syndrome CoV (MK651076), subgenus Pedacovirus porcine epidemic diarrhea virus (MK841495), subgenus Soravirus Shrew coronavirus isolate Shrew-CoV/Tibet2014 (NC_046955, 1 isolate); subgenus Sunacovirus Wencheng Sm shrew coronavirus isolate Xingguo-74 (NC_048211, 10 isolates); subgenus Luchacovirus Rodent coronavirus isolate RtMruf-CoV-1/JL2014 (KY370045, 15 isolates), Lucheng Rn rat coronavirus isolate Lijiang-71 (MT820627, 13 isolates), Lucheng Rn rat coronavirus isolate Lucheng-19 (NC_032730, 14 isolates), subgenus Minacovirus Mink coronavirus strain WD1127 (NC_023760, 552 isolates), Ferret coronavirus isolate FRCoV-NL-2010 (NC_030292, 10 isolates), Ferret coronavirus (LC215871, 9 isolates), subgenus Robacovirus BtRf-AlphaCoV/YN2012 (NC_0268824, 73 isolates), Rhinolophus affinis bat CoV HKU2-related isolate 160660 (MN611522, 75 isolates), Porcine enteric alphacoronavirus strain PEAV-GD-CH/2017 (MG742313, 347 isolates), Rhinolophus bat coronavirus HKU2 (NC_009988); subgenus Myotacovirus Bat alphacoronavirus isolate AMA_L_F (MT862548, 213 isolates), BtMr-AlphaCoV/SAX2011 (NC_028811, 789 isolates); subgenus Setracovirus NL63-related bat coronavirus strain BtKYNL63-15 ((KY073746, 266 isolates), subgenus Colacovirus Bat coronavirus CDPHE15/USA/2006 (NC_022103, 10 isolates); subgenus Decacovirus BtRf-AlphaCoV/HuB2013 (NC_028814, 33 isolates), Rousettus bat coronavirus HKU10 (NC_018871, 951 isolates), Hipposideros bat coronavirus HKU10 isolate TLC1343A (JQ989272, 937 isolates), Hipposideros pomona bat coronavirus HKU10-related isolate 160942 (MN611523, 947 isolates); subgenus Minunacovirus Miniopterus bat coronavirus HKU8 (NC_010438, 579 isolates), Bat coronavirus 1B strain AFCD307 (EU420137, 778 isolates); subgenus Nyctacovirus Alphacoronavirus Bat-CoV/P.kuhlii/Italy/206645-41/2011 (MH938448, 644 isolates), Alphacoronavirus sp. isolate WA2028 (MK472068, 615 isolates); β-CoVs subgenus Embecovirus murine hepatitis virus (MHV; NC_048217; AF208067), rat CoV Parker (NC_006213), rabbit CoV (JN874562), and bovine CoV (BCoV, U00735 and NC_003045; in this case except for sequences in related CoVs like hCoV-OC43 in Figure 8), and subgenus Hibecovirus Bat-Hp-betacoronavirus/ZHeijang 2013 (KF636752 and NC_025217) and Zaria bat CoV strain ZBCoV (HQ166910); δ-CoVs subgenus Buldecovirus Porcine deltacoronavirus (USA/Ohio444/2014, KR265862; MN942260); Common-moorhen CoV HKU21 (NC_016996); Night-heron CoV HKU19 (NC_016994); Munia CoV HKU13-3514 (NC_011550); Bulbul CoV HKU11-934 ([NC_011547](https://www.ebi.ac.uk/ena/data/view/NC_011547.1)); White-eye CoV HKU16 (NC_016991); Wigeon CoV HKU20 (NC_016995); Sparrow CoV HKU17 strain HKU17-6124 (JQ065045); Sparrow deltacoronavirus strain ISU73347 (MG812378, 176 isolates); and Thrush CoV HKU12-600 (FJ376621), subgenus Andecovirus Wigeon coronavirus HKU20 (NC_0169955, 4 isolates), subgenus Herdecovirus Night-heron coronavirus HKU19 (NC_016994, 1 isolate); and γ-CoVs subgenus Igavirus Infectious avian bronchitis virus (IABV; NC_001452; AY319651) and Turkey CoV (NC_010800), subgenus Cegacovirus Beluga Whale CoV SW1 (NC_010646) and Bottlenose dolphin CoV HKU22 isolate CF090327 (KF793825), and subgenus Brangacovirus Canada goose coronavirus strain Cambridge_Bay_2017 (NC_046965, 552 isolates).
